# High-throughput thermodynamic fingerprinting of protein–ligand interactions by DNA-directed focal molography

**DOI:** 10.64898/2026.07.03.736402

**Authors:** John Oehninger, Simona Notova, Andreas Frutiger

## Abstract

Thermodynamic characterization of biomolecular interactions is essential for understanding the enthalpic and entropic driving forces of molecular recognition, but established label-free techniques are limited either by bulk refractive-index sensitivity or by the lengthy thermal equilibration required to suppress it. Here, we used focal molography to investigate the temperature-dependent binding of the protein kinase A regulatory subunit (PKA-R) to cyclic AMP (cAMP) derivatives and to derive apparent thermodynamic signatures from kinetic measurements. We first validated the diffractometric readout under conditions that challenge refractometric sensors: the coherent mass density channel strongly suppressed temperature-induced bulk refractive-index effects and resolved binding in 50% human serum despite measurable non-specific adsorption, reducing the need for lengthy equilibration and buffer matching. We then combined focal molography with DNA-directed immobilization (DDI), allowing five cAMP derivatives to be presented in parallel on the same multiplexed chip and followed across five temperatures. This format yielded distinct, internally consistent apparent thermodynamic fingerprints for each derivative, separating ligands with similar affinities by their enthalpic and entropic contributions. Together, these results establish focal molography with DDI as a multiplexed workflow for comparative thermodynamic fingerprinting of biomolecular interactions at higher throughput.

## Introduction

The enthalpic and entropic decomposition of binding free energy provides mechanistic insight into the forces that govern molecular recognition and is therefore central to thermodynamic profiling in drug discovery and chemical biology [1]. In structure– activity relationship studies and lead-optimization campaigns, knowing whether closely related ligands achieve their affinity through enthalpic or entropic driving forces—and how those contributions track with chemical modification—can guide compound design beyond what affinity alone reveals. Realizing this potential requires methods that can deliver thermodynamic decompositions comparatively across many ligands under closely matched conditions, with sufficient throughput to support iterative chemistry.

Conventional biophysical techniques face distinct limitations in this setting. Surface plasmon resonance (SPR) can extract thermodynamic parameters from temperature-dependent kinetics via van ‘t Hoff and Eyring analyses, but is highly sensitive to bulk refractive-index changes and temperature drift; temperature ramping on SPR instruments therefore requires lengthy equilibration (30–60 min per step), making multi-temperature thermodynamic measurements low-throughput [2, 3]. Isothermal titration calorimetry (ITC) is the gold standard for solution-phase thermodynamics and is less affected by surface effects, but it requires large sample volumes and high protein concentrations, and each ligand must be titrated separately. Neither technique naturally combines higher-throughput kinetic characterization with multiplexed comparative thermodynamic profiling across multiple ligands. A second practical limitation is matrix compatibility. In complex biological matrices such as serum—where competing biomolecules, macromolecular crowding, and matrix viscosity can influence binding [4–7]— robust operation remains challenging for refractometric platforms because of bulk and non-specific adsorption artifacts, and for ITC because of large composition-dependent heat backgrounds.

Focal molography is a label-free optical method designed for biomolecular interaction analysis with reduced susceptibility to bulk and matrix artifacts. It employs a two-dimensional sub-micron pattern of molecular recognition sites in the shape of a diffractive lens, termed a *mologram* [8–11]. The pattern is composed of almost a thousand 200 nm ridges (binding sites) and referencing areas (grooves) in between (Fig 1A). Light diffracted by molecules bound to the ridges interferes constructively at a focal point, while contributions from the grooves cancel destructively. Specific analyte binding therefore increases the coherent mass contributing to this focal spot. The corresponding intensity increase is the diffractometric signal, reported as coherent mass density (CMD) [12]. Crucially, CMD preferentially reports mass bound in spatial register with the grating; uniformly distributed adsorption and homogeneous refractiveindex changes are strongly suppressed rather than fully absent from the experiment. Non-specific adsorption and bulk refractive-index changes, by contrast, affect ridges and grooves more similarly than patterned specific binding; they primarily shift the *position* of the focal spot while contributing little to its intensity (Fig 1B). This positional shift constitutes a built-in refractometric channel, reported as mass density (MD), which—like SPR—responds to all refractive-index changes regardless of their origin: specific binding, non-specific adsorption, and bulk temperature effects alike. The same instrument thus provides both channels simultaneously (Fig 1C): CMD for coherently patterned bound mass, MD for the total refractive-index response. Beyond the present model system, focal molography has been applied to affinity and kinetic characterization of proteins in complex matrices such as serum [13], to multiplexed affinity profiling of DNA-conjugated compound libraries [14], and to the real-time quantification of protein interactions in living cells [15–17].

**Fig 1.**
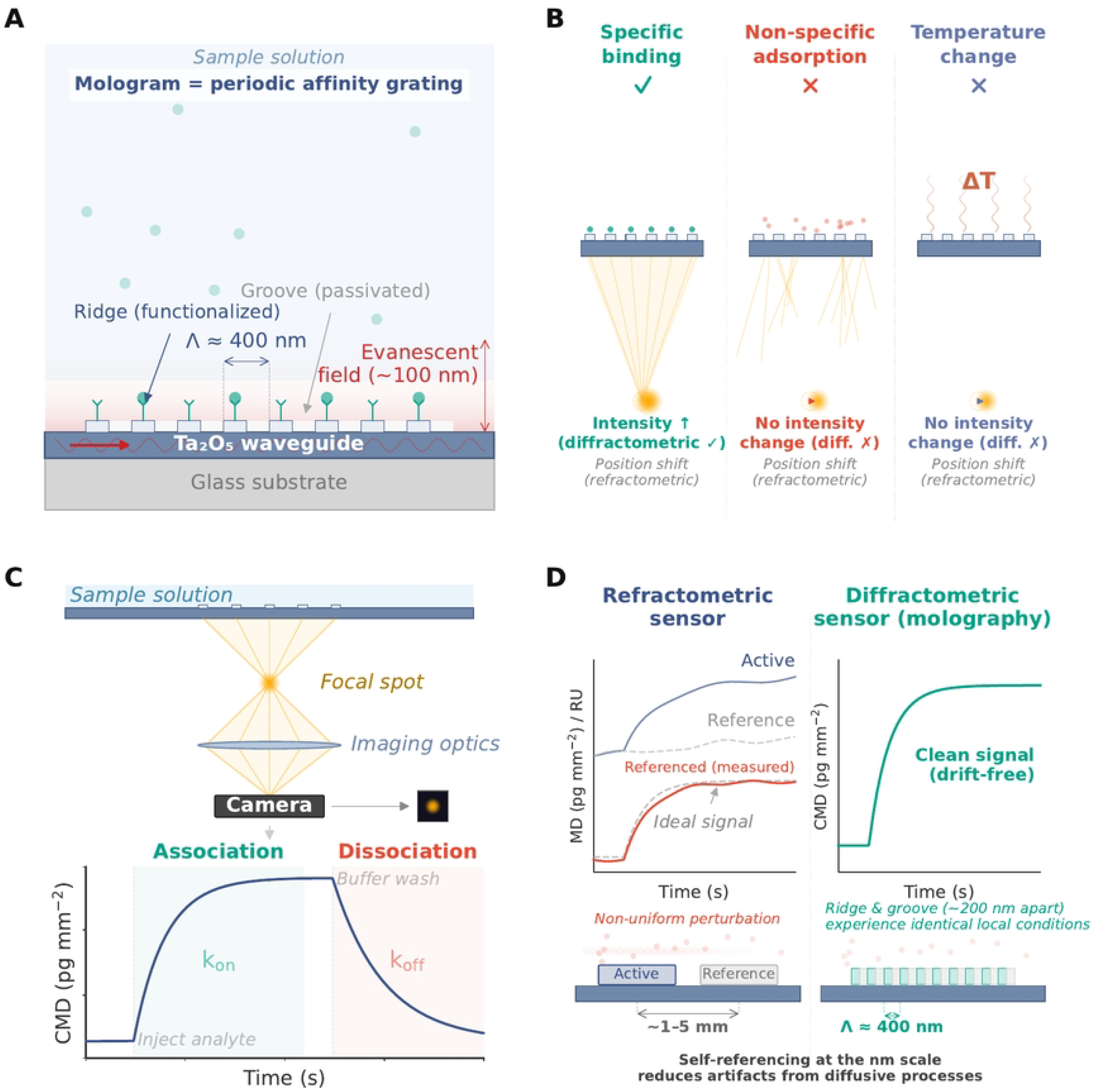
Principle of focal molography and self-referencing diffractometric detection. **(A)** Sensor cross-section. Laser light propagates as a guided mode in a Ta_2_O_5_ waveguide and generates an evanescent field extending ~100 nm above the surface. A mologram—a periodic affinity grating with period Λ ≈400 nm—consists of alternating functionalized ridges (carrying binding probes) and passivated grooves. **(B)** Coherent detection principle. Specific analyte binding on the ridges (left) increases the coherent mass in register with the grating period, producing a bright focal spot whose intensity rises (diffractometric signal). Spatially incoherent non-specific adsorption (center) and homogeneous bulk refractive-index changes such as temperature fluctuations (right) primarily shift the focal spot position (refractometric signal) and contribute little to its intensity when they affect ridges and grooves similarly. **(C)** Signal readout. A camera records the focal spot; its intensity yields the coherent mass density (CMD, pg mm^*−*2^) in real time. A representative sensorgram shows baseline, analyte association (*k*_on_) and dissociation (*k*_off_) phases. **(D)** Advantage of self-referencing at the sub-micron scale. In a conventional refractometric sensor (left), active and reference spots are separated by millimeters; spatially non-uniform perturbations (e.g. diffusing serum components) produce residual drift artifacts after subtraction, so the Referenced (measured) trace diverges from the Ideal signal (the true specific-binding contribution). In focal molography (right), ridges and grooves are interdigitated at ~200 nm spacing, so both experience closely matched local conditions—yielding a stable diffractometric signal with reduced dependence on a spatially separated reference. (Λ ≈400 nm denotes the grating period, i.e. the center-to-center pitch of one ridge–groove pair; it is not the optical wavelength.)

The key distinction from conventional refractometric techniques lies in this distributed, sub-micron-scale self-referencing mechanism [18, 19]: spatially homogeneous bulk contributions and much of the incoherent non-specific background cancel within the diffractometric channel itself, reducing the need for a spatially separated reference (Fig 1D). In contrast, SPR relies on a reference channel often millimeters away, where it becomes increasingly difficult to ensure that perturbations are correlated across signal and reference regions [20]. By suppressing homogeneous bulk contributions and much of the incoherent non-specific background, CMD should be less sensitive to temperature ramps and matrix-induced refractive-index artifacts than conventional refractometric readouts. Since the experiment follows surface-bound mass optically rather than measuring heat directly, it also avoids the large composition-dependent heat backgrounds that can complicate calorimetry in complex media. Together, these properties suggested that focal molography could provide a practical route to temperature-dependent kinetic measurements and, from them, apparent thermodynamic fingerprints by van ‘t Hoff and Eyring analyses.

In this study, we test this possibility using the interaction between the cAMP binding domain A of the bovine regulatory subunit RI*α* of PKA (bRI*α* 92–260, hereafter PKA-R) and cyclic AMP derivatives as a well-characterized model system in which cAMP binding to the regulatory subunit releases inhibition of the PKA catalytic subunit [21–23]. We begin by comparing the diffractometric and refractometric detection channels of the same instrument side by side, showing that the diffractometric channel remains stable during temperature ramps and in serum, whereas the refractometric channel exhibits large bulk artifacts. We then establish that focal molography resolves PKA-R binding kinetics for 6-AH-cAMP and 8-AHA-cAMP in both buffer and 50% human serum at 20°C with comparable kinetic parameters across media. To enable comparative thermodynamic profiling, we implement DNA-directed immobilization (DDI) via Watson–Crick hybridization, which allows simultaneous measurement of multiple ligands on the same multiplexed chip and yields distinct thermodynamic fingerprints for five cAMP derivatives across five temperatures. Together, these findings establish focal molography combined with DDI as a multiplexed platform for comparative thermodynamic fingerprinting at higher throughput.

## Materials and methods

### Reagents, proteins, and cAMP derivatives

Five cyclic AMP analogs carrying primary amine linkers were purchased from BIOLOG Life Science Institute (Bremen, Germany) and are shown in S1 Fig: N^6^-(6-aminohexyl)adenosine-3’,5’-cyclic monophosphate (6-AH-cAMP; CAS 66311-09-9, *M*_*W*_ 428.2 Da, cat. no. A031, lot 002002), 8-(6-aminohexylamino)adenosine-3’,5’-cyclic monophosphate (8-AHA-cAMP; CAS 39824-30-1, *M*_*W*_ 443.3 Da, cat. no. A011, lot 001004), 2-(6-aminohexylamino)adenosine-3’,5’-cyclic monophosphate (2-AHA-cAMP; CAS 214276-80-9, *M*_*W*_ 443.4 Da, cat. no. A053, lot 002007), 8-(2-aminoethyl)aminoadenosine-3’,5’-cyclic monophosphate (8-AEA-cAMP; CAS 61363-29-9, *M*_*W*_ 387.3 Da, cat. no. A016, lot 001010), and 8-(8-amino-3,6-dioxaoctylamino)adenosine-3’,5’-cyclic monophosphate (8-ADOA-cAMP; CAS 214272-05-6, *M*_*W*_ 475.4 Da, cat. no. A033, lot 001003). All analogs were dissolved in dimethyl sulfoxide (DMSO) at stock concentrations of 2.5– 10 mM and stored at −20°C. Of the five derivatives, 6-AH-cAMP and 8-AHA-cAMP were characterized at 20°C in the covalent format in both buffer A and 50% human serum (single-temperature kinetic comparison) and additionally in the DDI format in buffer; the remaining three (2-AHA-cAMP, 8-AEA-cAMP, 8-ADOA-cAMP) were profiled by DDI in buffer only. Temperature-dependent thermodynamic profiling (15–35°C) was performed exclusively in the DDI format in buffer.

The recombinant bovine RI*α* 92-260 protein (19,035.5 Da), a truncated form of the regulatory subunit of cAMP-dependent protein kinase A (PKA-R), was used in all experiments. This construct retains the essential cAMP-binding domain (residues 92–260), facilitating 1:1 stoichiometry with cAMP derivatives while simplifying data analysis compared to the full-length multi-domain protein, and is hereafter referred to as PKA-R [23]. The cDNA encoding RI*α* 92-260 was expressed in *E. coli* BL21 (DE3) RIL cells, and the protein was purified to obtain a nucleotide-free state following the procedure described in Moll et al. (2007) [21] and Hahnefeld et al. (2005) [22]. Briefly, the protein was eluted from 6-AH-cAMP agarose using 8 M urea in 20 mM MOPS (pH 7.0), followed by extensive dialysis against 150 mM NaCl, 20 mM MOPS, 10 mM MgCl_2_, and 1 mM ATP (pH 7.0). Protein purity was confirmed by SDS-PAGE, and its biological activity was verified using a phosphotransferase assay with Kemptide as substrate. Protein concentration was determined using a Bradford assay. All chemicals and reagents used were of analytical grade or higher.

### Instrumentation

Preliminary experiments and initial kinetic characterization in buffer and human serum were performed on a Callisto Pre-series prototype instrument (lino Biotech AG); a full description of that instrument and the associated data are provided in the Supporting Information (SI Methods, S6 Figs (a)–(c), SI Tables S2–S3).

All thermodynamic profiling experiments reported in this study were performed on the MACS^®^ Matchmaker (lino Biotech AG, Adliswil, Zurich, Switzerland; commercialized by Miltenyi Biotec GmbH, Bergisch Gladbach, Germany), the commercial successor instrument based on the same focal molography principle (S2 Fig). The MACS^®^ Matchmaker uses the same sensor chip format (9 mm ×18 mm) and mologram size (400 µm ×400 µm) as the Callisto, but features an 8×8 grid of 64 molographic measurement spots, with a within-row spacing of 785 µm and a between-row spacing of 900 µm. The flow chamber has a width of 2.1 mm, a height of 100 µm, and a length of 16 mm from inlet to outlet, and spans two rows of molograms (S2 Fig (b)).

### Sensor chip preparation, conjugation and immobilization

Sensor chip preparation involved three key steps: fabrication of molograms via reactive immersion lithography (RIL), chemical conjugation of cAMP derivatives to linker molecules, and subsequent immobilization of the conjugates onto the sensor surface.

### Mologram fabrication via reactive immersion lithography (RIL)

Each sensor chip features 64 molograms fabricated via RIL [9, 24]. This process uses a dielectric waveguide surface coated with photoactivatable polyacrylamide-grafted polyethylene glycol copolymer brushes (PAA-g-PEG-NH-PySNPPOC, SuSoS AG, Switzerland).

The brush layer forms a non-fouling background that suppresses non-specific binding while preserving functionality of the immobilized biomolecules [9].

A slight variation to the published protocol for the photolithography was employed: a more hydrophilic pyridine group was used as the photoprotective moiety of the primary amines (PySNPPOC – Carbamic acid 2-[4-ethyl-2-nitro-5-(pyridin-2-ylsulfanyl)phenyl]propyl ester] purchased from Orgentis Chemicals, Germany). Molograms were patterned by photolithography using a phase mask, which selectively removed the protective groups in the ridges, creating a line pattern of exposed primary amino groups. These amines were functionalized with 1 mM MeTz-PEG4-NHS (Methyltetrazine-(polyethylene)-N-hydroxysuccinimide, Vector Laboratories, USA) in HBS-T buffer (50 mM HEPES, 150 mM NaCl, pH 8.0) for 1 hour, then washed sequentially with DMSO, isopropanol (IPA), and Milli-Q water.

Following a flood exposure step to deprotect the remaining regions [10], the grooves were passivated with 10 mM MeO-PEG2-NHS (Broadpharm, USA) in HBS-T (pH 8.0). This resulted in spatially separated regions containing clickable MeTz-groups and PEG-passivated amines, embedded in a two-dimensional layer of the underlying PAA-g-PEG polymer roughly 10–20 nm in thickness in the hydrated state. Mologram architectures are denoted as [ridges|grooves]; the resulting structures are therefore referred to as [MeTz|PEG] molograms.

### Conjugation of cAMP derivatives

To enable site-specific immobilization, the cAMP derivatives were conjugated to TCO-PEG4-TFP linkers (Transcyclooctene-polyethyleneglycol-tetrafluorophenyl, Vector Laboratories, USA) via amine-reactive coupling. A 5:1 molar excess of cAMP derivative to TCO linker was used to drive the reaction to completion and avoid the need for purification.

Specifically, 200 µL of a 5000 µM cAMP derivative solution in DMSO was mixed with 2 µL of 100 mM TCO-PEG4-TFP (in DMSO) and 198 µL of HBS-T buffer (pH 8.0). The reactants were incubated for 1 hour at 15°C on a shaker at 600 rpm to ensure efficient conjugation between the TFP esters and primary amines on the cAMP derivatives.

### Ligand immobilization onto sensor surface (covalent format)

The resulting cAMP-TCO conjugates were applied directly to the sensor surface for immobilization via MeTz-TCO click chemistry [25]. A 100 µL aliquot of the conjugate solution was pipetted onto the chip and incubated for 1 hour at room temperature.

After incubation, the chip was sequentially washed in NaOH, toluene, isopropanol, Milli-Q water, and 3 M guanidine hydrochloride (GuHCl), with each wash lasting 30 seconds on a shaker. The chip was then dried with nitrogen gas and stored at 4°C in an Eppendorf tube until use.

Immediately before each experiment, the chip was rinsed with Milli-Q water, dried with inert gas, and mounted into the flow chamber for measurements.

### DNA-directed immobilization workflow

#### Conjugation of cAMP derivatives to TCO-functionalized oligonucleotides

For DNA-directed immobilization, each cAMP derivative was conjugated to a TCO-functionalized oligonucleotide (TCO-oligo, Biomers, Germany) via a two-step procedure. In the first step, the primary amine on the cAMP derivative was reacted with a methyltetrazine (MeTz) linker using NHS ester chemistry. To this end, a semi-stock of each cAMP derivative was prepared at 100 µM in 150 µL PBS (pH 8.0), and the MeTz-PEG4-NHS linker (Methyltetrazine-PEG4-NHS Ester, CCT-1069-100, Vector Laboratories, USA; *M*_*W*_ = 533.53 Da) was prepared at 100 µM by diluting 1 µL of 100 mM DMSO stock into 999 µL PBS (pH 8.0). For each cAMP derivative, 150 µL (15,000 pmol) of cAMP semi-stock were combined with 75 µL (7,500 pmol) of linker semi-stock (molar ratio 2:1, cAMP:linker) and incubated for 1 h at 25°C at 800 rpm.

In the second step, TCO-functionalized oligonucleotides (20-mer, TCO on the 5′ end) were added at 5,000 pmol per reaction (333 µL from a 15 µM stock, corresponding to 31.5 µg) and incubated for an additional 1 h at 25°C at 800 rpm. The conjugation relies on the inverse electron demand DielsAlder (IEDDA) click reaction between the TCO group on the oligonucleotide and the MeTz group on the cAMP conjugate. The incubation was extended relative to the covalent format to account for the lower effective oligo concentration and to ensure complete MeTzTCO reaction. Each cAMP derivative was paired with a designated complementary oligonucleotide strand: 8-ADOA-cAMP with cs01, 2-AHA-cAMP with cs02, 6-AH-cAMP with cs04, 8-AEA-cAMP with cs06, and 8-AHA-cAMP with cs07.

#### Purification by ion-exchange chromatography

The resulting cAMPoligonucleotide conjugates were purified by ion-exchange chromatography (IEX) on an Ä KTA system using a Resource Q column, following the standard IEX Resource Q method with a modified elution flow rate of 0.5 mL/min. A 500 µL sample loop was used and 500 µL of the reaction mixture were loaded per run. Conjugate-containing fractions were identified, pooled, and quantified by NanoDrop absorbance measurement. Conjugation yields and purification outcomes for all five derivatives are summarized in Table 1.

**Table 1.**
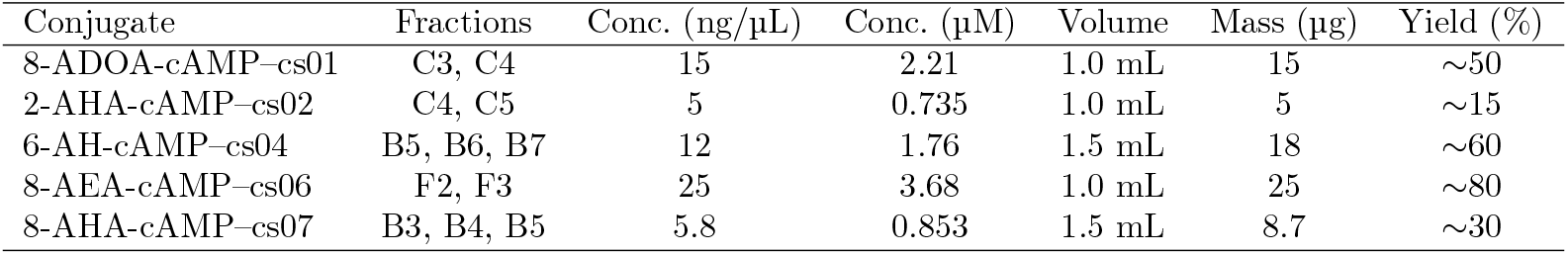
Purification fractions and conjugation yields for all cAMP–oligonucleotide conjugates. Concentration and mass were determined by NanoDrop absorbance. Yield is expressed relative to the input TCO-oligo (5,000 pmol, 31.5 µg).

Functional quality control of all purified conjugates was performed directly on the focal molography instrument to confirm capture activity prior to use in thermodynamic profiling experiments.

### Experimental setup and procedure

All interaction analyses were conducted in a MOPS-based buffer composed of 150 mM NaCl, 20 mM MOPS, 10 mM MgCl_2_, 15 µM BSA and 0.05% (v/v) Tween-20 at pH 7.0, termed *buffer A* throughout. BSA was included as a blocking agent to reduce non-specific adsorption of PKA-R to the sensor surface.

For experiments conducted in human serum on the MACS^®^ Matchmaker, normal human serum (MP Biomedicals, cat. no. 2930149, lot U1123331304; ordered via Fisher Scientific, cat. no. 11475045; storage at −20°C) was diluted to 50% in *buffer A. Buffer A* was not refractive-index matched to the human serum preparation. This preparation exhibited measurable non-specific binding to covalently functionalized sensor chips, which is addressed in the relevant results sections. A thyroid-stimulating hormone (TSH)-depleted serum preparation (lot HD2403001) used in prototype instrument experiments is described in the Supporting Information (SI Methods).

For buffer measurements, PKA-R was diluted in *buffer A* during association phases. For serum measurements, PKA-R was diluted in 50% human serum during association, whereas dissociation phases and buffer-wash steps used *buffer A*. Thus, buffer and serum experiments followed the same injection sequence except for the matrix used during association. The standardized injection sequences for multi-cycle kinetics (MCK) experiments on the MACS^®^ Matchmaker and for DNA-directed immobilization single-cycle kinetics (DDI-SCK) experiments are summarized in Tables 2 and 3, respectively. For MCK experiments, regeneration was performed using three successive injections of regeneration solution (3 M GuHCl and 125 mM NaOH) to ensure complete removal of bound analyte between cycles.

**Table 2.** Injection sequence for multi-cycle kinetics (MCK) experiments on the MACS^®^ Matchmaker.

| Type | Sample | Duration | Flow rate |
| --- | --- | --- | --- |
| Baseline | Buffer A | ~2 min | 150 µL/min |
| Association | PKA-R | 72 s | 150 µL/min |
| Dissociation | Buffer A | ~4.5 min | 400 µL/min |
| Regeneration | Regeneration solution | 3 × ~2.5 min | 300 µL/min |
| Rinse | Buffer A | ~1 min | 300 µL/min |

**Table 3.** Injection sequence for DNA-directed immobilization single-cycle kinetics (DDI-SCK) experiments on the MACS^®^ Matchmaker. The SCK block consists of a reference injection at 0 nM followed by nine ascending PKA-R concentrations (2.5, 5, 10, 20, 40, 80, 160, 320, 640 nM), for a total of ten injection steps. Each association/dissociation step is repeated for every concentration without regeneration between injections. For kinetic fitting, the 0 nM blank step and the 640 nM step are excluded (see Methods), leaving eight fitted concentrations (2.5–320 nM).

| Type | Sample | Duration | Flow rate |
| --- | --- | --- | --- |
| Pre-regeneration | Regeneration solution | 60 s | 300 µL/min |
| Immobilization baseline | Buffer A | ~2 min | 300 µL/min |
| Immobilization | cAMP–oligo mix (200 nM each) | 5 min | 10 µL/min |
| Equilibration | Buffer A | ~5 min | 300 µL/min |
| Association (×10) | PKA-R (0–640 nM) | 72 s per step | 150 µL/min |
| Dissociation (10) | Buffer A | ~4.5 min per step | 400 µL/min |
| Post-regeneration | Regeneration solution | 60 s | 300 µL/min |

The sensor chip was mounted in the aforementioned three-channel flow chamber. Temperature control was maintained at 15°C, 20°C, 25°C, 30°C and 35°C using the Thorlabs LK220 Thermoelectric Liquid Chiller. Sixteen molograms were recorded simultaneously per flow channel.

#### Temperature ramp validation

To validate the insensitivity of the diffractometric detection to temperature-induced bulk refractive-index changes, a continuous 25–45–25 °C temperature ramp was performed on the MACS^®^ Matchmaker in buffer only (no analyte). The ramp lasted approximately 100 min and was recorded on all 16 molograms simultaneously (full 224 min recording in S3 Fig.). The coherent mass density (CMD, diffractometric channel) and the refractometric channel were monitored in parallel. For the refractometric channel, both unreferenced single-sensor traces and referenced traces (consecutive sensor-pair subtraction) were compared to assess the effectiveness of active referencing against thermal drift.

#### Covalent format — single-temperature kinetic comparison

For the covalent immobilization format, MCK titration series were acquired at 20°C only, in order to compare the kinetic behavior of 6-AH-cAMP and 8-AHA-cAMP in buffer A and 50% human serum. At each condition, a full MCK titration was performed with GuHCl/NaOH regeneration between individual analyte concentration cycles as described in Table 2. Between the serum and buffer titrations on the same chip, an enzymatic Trypsin-EDTA regeneration was applied to remove non-specifically adsorbed serum material; the protocol and a representative sensorgram are provided in SI Section S8 (S8 Fig). Temperature-dependent profiling on the covalent format is not reported in this study.

#### DDI format — measurement order

For the DNA-directed immobilization (DDI) format, all five cAMP–oligonucleotide conjugates were immobilized simultaneously from a single premixed solution (200 nM each), enabling multiplexed kinetic profiling in a single SCK cycle. Each conjugate hybridized to its designated complementary capture oligo on the chip, so that one SCK run yielded binding data for all five compounds in parallel.

DDI-SCK data were acquired in repeated immobilize–measure–regenerate cycles across several experimental sessions covering the same five temperatures (15, 20, 25, 30, 35°C). Most single-temperature sessions contained three SCK cycles, while one experiment measured 25, 15 and 30°C sequentially. After excluding SCK phases affected by air-bubble or other visible acquisition artifacts, the final analyzed dataset contained 5, 9, 7, 5 and 6 included SCK cycles at 15, 20, 25, 30 and 35°C, respectively. Because each active flow channel spans two mologram rows, each included cycle contributed two positive molograms per compound. This yielded *N* = 10, 18, 14, 10 and 12 individual fits per compound at 15, 20, 25, 30 and 35°C, respectively (64 fits per compound in total). Three negative-control oligo identities without a cAMP conjugate (cs03, cs05, cs08) were present in the same flow channel; these showed no detectable PKA-R binding during the SCK phase (S4 Fig (c)), supporting that the fitted kinetic signal originates from cAMP–PKA-R binding at the cognate mologram spots. Negative-control molograms were excluded from kinetic analysis.

### Data acquisition and kinetic analysis

Data were collected at the maximum available sampling frequency of the MACS^®^ Matchmaker (approximately 1.5 Hz). The instrument generated sensorgrams that were subsequently analyzed to extract kinetic and thermodynamic information using a dedicated kinetic fitting library provided by lino Biotech AG. This code leverages the widely-used Python module `lmfit` to perform nonlinear least-squares fits [26]. To analyze the binding data, a Langmuir 1:1 binding model was employed for the majority of compounds. This model assumes a simple bimolecular interaction and fits three parameters *k*_on_, *k*_off_ and *R*_max_. For two DDI compounds—2-AHA-cAMP and 8-ADOA-cAMP—the standard Langmuir 1:1 model was extended with a non-dissociating fraction parameter *f*_nd_ (Langmuir 1:1 NonDissRelative model). In this variant the association phase follows standard 1:1 kinetics, but during dissociation a fraction *f*_nd_ of the bound analyte remains permanently associated while the remainder (1 *− f*_nd_) dissociates with rate constant *k*_off_, yielding four fitted parameters (*k*_on_, *k*_off_, *R*_max_, *f*_nd_). Specifically, the dissociation-phase response is modeled as:

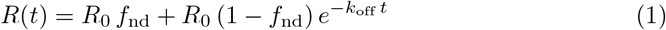

where *R*_0_ is the response at the onset of dissociation. The kinetic parameters extracted include the association rate constant (*k*_on_), dissociation rate constant (*k*_off_), and equilibrium dissociation constant (*K*_D_). The equilibrium dissociation constant was calculated as:

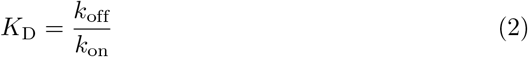

The fitting process was conducted globally, fitting all data points across different analyte concentrations simultaneously to enhance robustness and reliability.

### MCK data preprocessing and fitting parameters

Prior to kinetic fitting, the CMD sensorgrams were preprocessed by subtracting the blank (0 nM analyte) injection cycle from all concentration cycles, yielding ΔCMD sensorgrams (blank-cycle-subtracted, unreferenced; distinct from the spatial self-referencing of the mologram grating). For human serum experiments, this blank-cycle subtraction removes the bulk serum response. For buffer experiments, it has no measurable effect but was applied uniformly to maintain an identical processing pipeline across conditions.

The fitting parameters differed between conditions to account for their distinct binding characteristics. For buffer A experiments, the highest analyte concentration (640 nM) was excluded from the global fit, and the dissociation phase was truncated at 200 s. The concentration exclusion was necessary because the 640 nM data deviated from ideal 1:1 Langmuir behavior, and the time cutoff avoided fitting the slow non-dissociating tail observed in buffer. For 50% human serum experiments, all nine analyte concentrations (2.5–640 nM) were included and the full dissociation phase was fitted without truncation. In all cases, 16 sensor spots were fitted individually per temperature and condition.

### DDI-SCK data preprocessing and fitting parameters

For DDI-SCK experiments, the 0 nM blank injection and its subsequent dissociation phase were removed from the data prior to fitting. Each CMD sensor trace was then offset to zero by subtracting the mean signal from a 30-second window immediately preceding the first non-blank association injection (a display offset only; not a blank-cycle subtraction). The highest analyte concentration (640 nM) was excluded from the global fit, as for MCK, leaving eight fitted concentrations (2.5–320 nM). The full dissociation phase was fitted without time truncation.

The standard Langmuir 1:1 model was used for 6-AH-cAMP, 8-AHA-cAMP and 8-AEA-cAMP, while 2-AHA-cAMP and 8-ADOA-cAMP were fitted with the Langmuir 1:1 NonDissRelative model (Eq. 1) to account for their non-dissociating fraction. Sensors associated with negative control oligonucleotides (cs03, cs05, cs08) were excluded from kinetic analysis.

Standard-state thermodynamic parameters, such as Gibbs free energy (Δ*G*°), enthalpy (Δ*H*°), and entropy (Δ*S*°), were derived from van’t Hoff plots using temperature-dependent binding data. For the equilibrium dissociation constant, parameters were defined relative to the standard-state concentration *C*° = 1 M, yielding the dimensionless ratio *K*_D_*/C*°. These parameters were calculated using the following relationships:

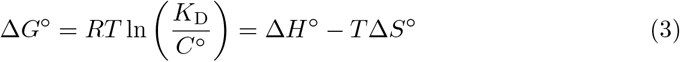

and

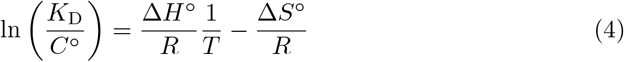

Here, *R* is the universal gas constant and *T* is the absolute temperature in Kelvin. With *C*° = 1 M and *K*_D_ expressed in M, this standard-state convention is numerically identical to using the fitted *K*_D_ values directly, but it defines the reported equilibrium entropy and free energy with respect to a 1 M standard state. The slope of the van’t Hoff plot (ln(*K*_D_*/C*°) vs. 1*/T*) provides the enthalpy change (Δ*H*°), while the intercept gives the entropy change (Δ*S*°).

### Calculation of activation parameters

The thermodynamic parameters of the transition state for the association 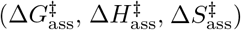 and dissociation 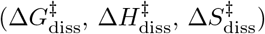 were calculated using kinetic rate constants (*k*_on_ and *k*_off_). Because *k*_on_ is a second-order rate constant (M^*−*1^s^*−*1^), association activation parameters were defined relative to the standard-state concentration *C*° = 1 M, converting *k*_on_ into the standard-state first-order rate constant *k*_on_*C*°. According to transition state theory, the association process is then described by:

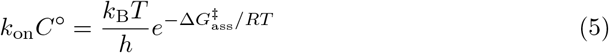

Here, *h* is the Planck constant and *k*_B_ the Boltzmann constant. With *C*° = 1 M and *k*_on_ expressed in M^*−*1^s^*−*1^, this standard-state convention is numerically identical to using the fitted *k*_on_ values directly, but it defines the reported association entropy and free energy of activation with respect to a 1 M standard state. After substituting 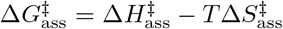 and taking the natural logarithm, this yields the linearized Eyring equation:

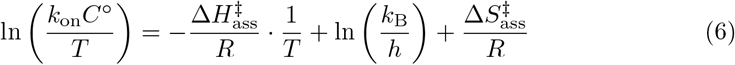

In this form, plotting ln(*k*_on_*C*°*/T*) against 1*/T* yields a line, where the slope is 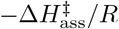 and the intercept is ln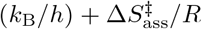. This allows the determination of both the enthalpy 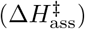 and entropy 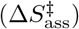 of activation. For dissociation, the same procedure applies directly to the first-order rate constant *k*_off_.

### Controls and validation

#### Positive control: verification of specific PKA-R binding

To validate the successful immobilization of cAMP derivatives on the sensor chip, a positive control experiment was performed by injecting 320 nM PKA-R in *buffer A* over the sensor surface immobilized with the 6-AH-cAMP analog. This concentration was chosen based on prior studies to generate a strong response [21]. As shown in S4 Fig (a)A, the injection of PKA-R resulted in a sharp and substantial increase in coherent mass density (≈20 pg/mm^2^ within 30 seconds), indicating a successful and specific binding event. This increase stabilized at a higher level, consistent with free PKA-R molecules interacting with the immobilized cAMP on the sensor surface. These results demonstrate detection of specific PKA-R–cAMP interactions and successful immobilization of cAMP derivatives.

#### Negative control: competition with free cAMP

To further assess the specificity of the measured interaction, a negative control experiment was conducted using a pre-formed PKA-R–cAMP complex. A solution containing 160 nM PKA-R and 1.6 µM 6-AH-cAMP (1:10 molar ratio) was injected over the sensor chip to occupy nearly all PKA-R binding sites before exposure to the immobilized cAMP. As illustrated in S4 Fig (a)B, the injection of the PKA-R–cAMP complex resulted in no significant change in coherent mass density, with only minor fluctuations within a range of ≈ 0.5 pg/mm^2^. These fluctuations are consistent with baseline noise and indicate that pre-saturated PKA-R did not produce a detectable non-specific CMD response under these conditions. The absence of a binding response supports the specificity of the assay and indicates that the observed binding signal is dominated by free, unbound PKA-R interacting with immobilized cAMP.

#### Assessment of non-specific binding in 50% human serum

Non-specific adsorption of serum components was characterized by blank injections of each serum preparation on chips covalently functionalized with 6-AH-cAMP prior to kinetic measurements. Normal human serum (lot U1123331304, 50% in *buffer A*) produced a measurable non-specific signal of approximately 16 pg mm^*−*2^ with partial recovery upon buffer washout, indicating residual adsorption onto the covalently functionalized chip surface that must be considered when interpreting serum binding data. In contrast, the TSH-depleted serum preparation used on the Callisto Pre-series reader produced no detectable non-specific signal, demonstrating that different serum preparations can differ substantially in their non-specific binding behavior (S4 Fig (b)).

### Reproducibility and statistical analysis

#### Reproducibility design

- Covalent format: kinetic measurements at 20°C in buffer and 50% human serum used 16 active sensors per condition; matrix comparison is based on this single-temperature dataset.
- DDI format: all five cAMP derivatives were profiled simultaneously in each included DDI-SCK cycle. After quality exclusions, 5, 9, 7, 5 and 6 cycles were retained at 15, 20, 25, 30 and 35°C, respectively, corresponding to *N* = 10, 18, 14, 10 and 12 individual fits per compound at these temperatures.

#### Kinetic parameter statistics

For covalent MCK measurements, 16 sensor spots were fitted individually per temperature and condition. For DDI-SCK measurements, each retained positive mologram/cycle trace was fitted individually, yielding per-fit estimates of *k*_on_, *k*_off_, and *K*_*D*_ together with the residual *χ*^2^ of each fit. In the DDI isoaffinity and thermodynamic scatter plots, individual retained fits are shown as points. For tabulated 25 °C DDI kinetic parameters, values are arithmetic mean ± sample standard deviation across the 14 retained individual fits at that temperature.

#### Thermodynamic parameter uncertainties

For DDI-SCK thermodynamic analysis, the retained individual fits were first summarized at each temperature as *χ*^2^-weighted means of *K*_*D*_, *k*_on_, and *k*_off_, using weights 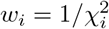 after clipping very small *χ*^2^ values. Central equilibrium and activation parameters (Δ*H*, Δ*S*, Δ*G*, Δ*H*^*‡*^, Δ*S*^*‡*^, Δ*G*^*‡*^) were then obtained from linear van ‘t Hoff and Eyring fits across the five temperature means; per-temperature regression weights were the sums of the corresponding individual-fit weights. The reported *N* for DDI-SCK (*N* = 10, 18, 14, 10 and 12 at 15, 20, 25, 30 and 35°C; total *N* = 64 per compound) denotes the number of retained individual kinetic fits contributing to these temperature means.

Uncertainties on the DDI-SCK thermodynamic parameters are bootstrap standard deviations. For each compound, individual fits were resampled with replacement within each temperature group, *χ*^2^-weighted temperature means were recomputed, and the resulting five resampled temperature means were refitted with ordinary linear van ‘t Hoff and Eyring regressions over 1,000 bootstrap iterations. Error bars on DDI thermodynamic bar charts and the uncertainties reported in the DDI thermodynamic tables correspond to the standard deviation of these bootstrap estimates. They therefore capture within-assay variability among retained molograms, cycles, and experimental sessions, but not independent chip-to-chip, protein-preparation, or conjugate-preparation variability.

## Results

We first evaluate the temperature stability and matrix tolerance of the diffractometric detection channel of focal molography — prerequisites for reliable thermodynamic profiling. We then establish baseline kinetics for 6-AH-cAMP and 8-AHA-cAMP binding to PKA-R in buffer and 50% human serum at a single temperature on the covalent format, showing that kinetic parameters remain comparable across media. Finally, we introduce DNA-directed immobilization (DDI) to enable multiplexed, temperature-dependent thermodynamic profiling of five cAMP derivatives simultaneously, yielding distinct enthalpic and entropic fingerprints for each ligand.

### Diffractometric focal molography is temperature-stable and matrix tolerant

Before thermodynamic profiling, we first demonstrate two key prerequisites for reliable temperature-dependent measurements: stability against bulk refractive-index changes caused by temperature and tolerance to complex biological matrices. As described in the Introduction (Fig 1B), the focal molography instrument provides two simultaneous detection channels: the diffractometric channel measures the intensity of the focal spot, which preferentially reports coherently patterned bound mass (CMD), while the refractometric channel measures the positional shift of the focal spot, reported as mass density (MD), which — like SPR — responds to any refractive-index change including bulk effects and non-specific adsorption. To put the magnitudes discussed below into perspective, a typical protein monolayer corresponds to approximately 400 pg mm^*−*2^ in CMD and 2000–4000 pg mm^*−*2^ in MD.

To quantify the temperature sensitivity of each detection channel, a continuous buffer-only temperature ramp from 25°C to 45°C and back to 25°C was recorded on the MACS^®^ Matchmaker (16 sensors per flow channel). Three channels were compared: CMD, the unreferenced mass density (MD, zero-offset to starting value), and the actively referenced mass density (MD_ref_) obtained by pairwise subtraction of consecutive sensors (Fig. 2A–C; full recording in S3 Fig.). CMD remains within a range of less than 1 pg mm^*−*2^ throughout the entire 20°C ramp — less than 0.25% of a protein monolayer — showing effective stability against temperature-induced bulk refractive-index changes. By contrast, MD shows a large apparent signal change of several thousand pg mm^*−*2^ — on the order of one to two refractometric monolayer equivalents. Active referencing by consecutive sensor-pair subtraction (MD_ref_) substantially suppresses this artifact, yet residual drift still spans up to ~ ± 1200 pg mm^*−*2^, on the order of a refractometric monolayer. Temperature ramping on refractometric biosensors such as SPR therefore requires lengthy equilibration periods — often 30 to 60 minutes per step — making thermodynamic measurements a low-throughput process. By reducing this equilibration requirement, focal molography enables faster and more practical determination of thermodynamic parameters.

**Fig 2.**
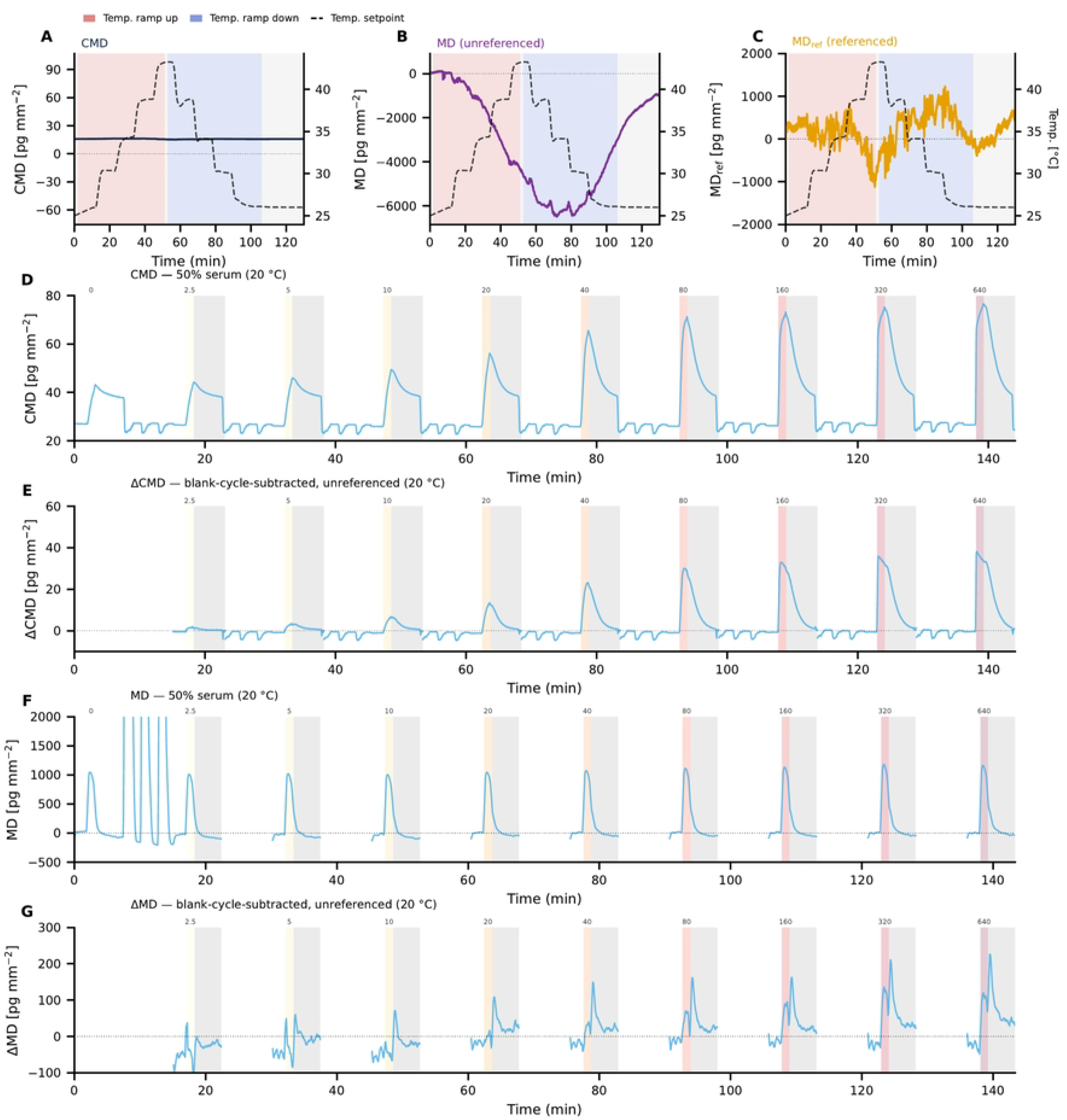
Diffractometric focal molography is temperature-stable and matrix-tolerant. Abbreviations used throughout: CMD, coherent mass density (diffractometric; preferentially reports coherently patterned bound mass); MD, mass density (refractometric; responds to all refractive-index changes); ΔCMD and ΔMD, blank-cycle-subtracted (unreferenced) traces (panels E and G); MD_ref_, actively sensor-pair-referenced MD (panel C). **Top row (A–C):** Buffer-only 25–45–25 °C temperature ramp on the MACS^®^ Matchmaker (~100 min ramp portion shown; full 224 min recording in S3 Fig.); red/blue shading: ramp-up/ramp-down; dashed line: heater/chiller temperature (right axis). **(A)** CMD: temperature-induced signal change below 1 pg mm^*−*2^ throughout, orders of magnitude below a typical protein monolayer (~400 pg mm^*−*2^). **(B)** MD (unreferenced; zero-offset to starting value): bulk refractive-index change produces several thousand pg mm^*−*2^ apparent signal. **(C)** MD_ref_ (consecutive sensor-pair subtraction): residual drift spans up to ~±1200 pg mm^*−*2^, on the order of an SPR monolayer (2000–4000 pg mm^*−*2^). **Bottom rows (D–G):** Multi-cycle kinetics of 6-AH-cAMP / PKA-R at 20 °C in 50% human serum on the MACS^®^ Matchmaker; concentrations 0–640 nM (log scale, color-coded); gray bands: dissociation windows. **(D)** CMD: binding responses clearly resolved without correction. **(E)** ΔCMD (blank-cycle-subtracted, unreferenced): clean sensorgram acquired in 50% human serum. **(F)** MD: large cycle-to-cycle nonspecific responses from serum almost completely mask binding. **(G)** ΔMD (blank-cycle-subtracted, unreferenced): binding responses become visible, but substantial artifacts remain.

The advantage of diffractometric detection is equally striking in complex biological matrices. MCK experiments of 6-AH-cAMP binding to PKA-R in 50% human serum show that CMD cleanly resolves all nine concentration cycles without any correction (Fig. 2D), and blank-cycle subtraction yields publication-quality ΔCMD sensorgrams (Fig. 2E). In stark contrast, MD is overwhelmed by non-specific serum signals that almost completely mask the binding responses (Fig. 2F). This panel also reveals the three regeneration injections of 3 M GuHCl with 125 mM NaOH between concentration cycles, which produce massive bulk refractive-index spikes of approximately 15,000 pg mm^*−*2^ — four to seven refractometric monolayer equivalents — in MD, while CMD in panel D shows virtually no response to these same injections. Even after blank-cycle subtraction, ΔMD retains substantial artifacts (Fig. 2G). Together, these results confirm that the diffractometric detection of focal molography is well suited for temperature-dependent kinetic measurements in complex media.

### MCK kinetics of 6-AH-cAMP and 8-AHA-cAMP at 20 °C in buffer and serum

Kinetic characterization of 6-AH-cAMP and 8-AHA-cAMP binding to PKA-R was first performed on the Callisto Pre-series prototype at 20 °C in both buffer A and 50% TSH-depleted human serum (S6 Figs (b)–(c), SI Tables S2–S3). These initial experiments showed that focal molography resolves specific PKA-R binding in serum with sensorgram shapes and signal amplitudes comparable to those obtained in buffer. Blank injection of the TSH-depleted serum produced no detectable non-specific signal in the CMD channel (*<*1 pg/mm^2^; S4 Fig (b)A), indicating low non-specific adsorption for this serum preparation under these conditions.

On the MACS^®^ Matchmaker, the same MCK protocol was performed at 20 °C with 16 sensors per flow channel, providing substantially better statistics than the single-sensor Callisto experiments. Representative full MCK traces for both analytes in buffer and serum are shown in Fig 3A–D. In buffer, the sensorgrams exhibit slow dissociation kinetics with evidence of a non-dissociating component at the highest concentrations; fits were therefore restricted to the initial 200 s of the dissociation phase and the 640 nM cycle was excluded (Fig 3E, F). In 50% human serum, the raw CMD again resolves concentration-dependent binding, and blank-subtracted sensorgrams yield well-behaved Langmuir 1:1 fits (Fig 3J, K). Box plots of the fitted kinetic parameters across all 16 sensors (Fig 3G–I, L–N) show closely overlapping distributions between buffer and serum for both analytes (Table 4); the corresponding per-sensor 1:1 Langmuir fits for all 16 sensors in each of the four conditions are shown in S5 Fig (a)–(d). The association rate *k*_on_ does not differ measurably between the two matrices, while *k*_off_ and *K*_D_ are marginally higher in serum (by up to ~20 %). A shift of this magnitude lies within the run-to-run reproducibility of surface-based kinetic assays and supports the conclusion that, under these conditions, the complex matrix did not materially alter the measured kinetics.

**Table 4.** Per-sensor MCK kinetic parameters for PKA-R binding to 6-AH-cAMP and 8-AHA-cAMP at 20 °C in buffer A and 50% human serum.

| Ligand | Matrix | $k_{\text{on}} (\times 10^5 \text{ M}^{-1} \text{ s}^{-1})$ | $k_{\text{off}} (\times 10^{-3} \text{ s}^{-1})$ | $K_D$ (nM) | $N$ |
| --- | --- | --- | --- | --- | --- |
| 6-AH-cAMP | Buffer A | $6.15 \pm 0.19$ | $10.11 \pm 0.26$ | $16.5 \pm 0.1$ | 16 |
| 6-AH-cAMP | 50% serum | $6.05 \pm 0.24$ | $11.89 \pm 0.44$ | $19.7 \pm 0.4$ | 16 |
| 8-AHA-cAMP | Buffer A | $5.59 \pm 0.16$ | $12.84 \pm 0.27$ | $23.1 \pm 0.4$ | 16 |
| 8-AHA-cAMP | 50% serum | $5.27 \pm 0.17$ | $14.45 \pm 0.38$ | $27.6 \pm 0.4$ | 16 |

**Fig 3.**
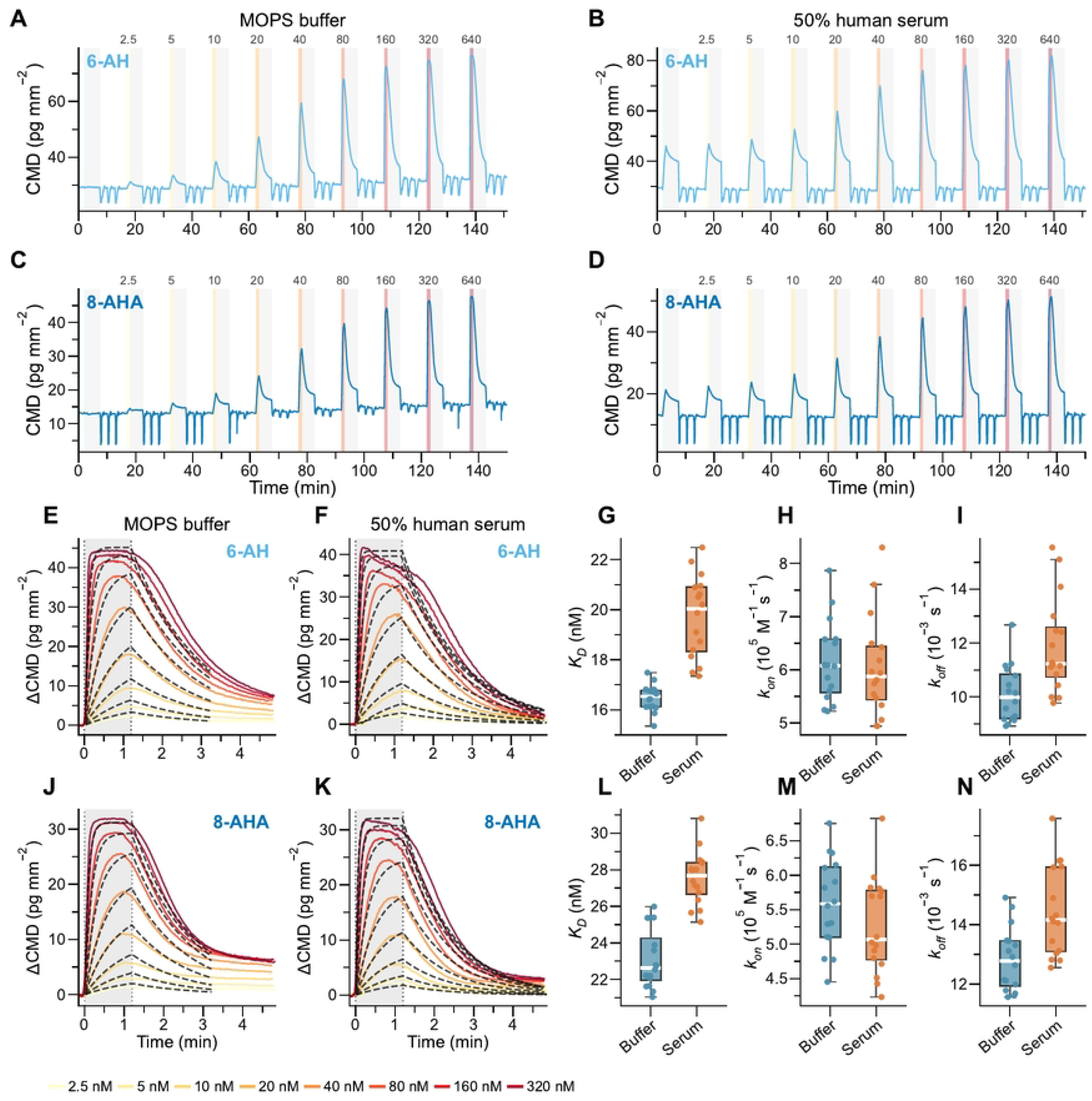
Multi-cycle kinetics of 6-AH-cAMP and 8-AHA-cAMP at 20 °C in buffer A and 50% human serum. **(A–D)** Full MCK experiment traces (~150 min) showing CMD (raw coherent mass density; self-referenced by the mologram grating, no additional baseline subtraction required) for nine successive PKA-R concentration steps (2.5–640 nM). Association phases are color-coded by concentration (yellow to red); dissociation phases are shaded gray. Concentration labels (nM) are shown above each association band. **(E–F, J–K)** Overlaid ΔCMD sensorgrams (blank-cycle-subtracted, unreferenced) for each concentration series. Coloured traces show experimental data; dashed black lines show individual stored Langmuir 1:1 fits (640 nM cycle excluded in buffer; dissociation truncated at 200 s in buffer to avoid fitting the non-dissociating tail). **(G–I, L–N)** Box plots (all 16 sensors, 20 °C) comparing buffer A (teal) and 50% human serum (amber) for the equilibrium dissociation constant *K*_*D*_, the association rate constant *k*_on_, and the dissociation rate constant *k*_off_. Individual sensor values are overlaid as jittered points. Distributions overlap closely between buffer and serum for both analytes; *k*_on_ does not differ measurably, whereas *k*_off_ and *K*_*D*_ are at most ~20 % higher in serum—within assay reproducibility and consistent with the same interaction (numerical values in Table 4).

Because the TSH-depleted serum was available only in limited quantity, the MACS^®^ Matchmaker MCK experiments used a normal human serum (lot U1123331304). Unlike the TSH-depleted preparation, this serum produced measurable non-specific adsorption of approximately 16 pg mm^*−*2^ on the covalently functionalized chips (S4 Fig (b)B). This difference highlights that non-specific binding is serum-preparation-specific and must be assessed independently for each lot and matrix.

### DNA-directed immobilization enables multiplexed parallel ligand presentation

To move from single-ligand kinetic characterization to comparative thermodynamic profiling across multiple compounds, we implemented DNA-directed immobilization (DDI). In this format, cAMP–oligonucleotide conjugates are presented on the sensor surface via Watson–Crick hybridization to complementary capture strands (Fig 4A, B). Compared with direct covalent attachment, DDI offers three properties relevant to comparative thermodynamic profiling: (i) multiple ligands self-sort simultaneously onto a single multiplexed chip in one injection step, so several compounds can be measured in parallel on the same chip under closely matched local conditions; (ii) ligand density and orientation are controlled by the capture-strand density and hybridization efficiency rather than by random surface chemistry; and (iii) the surface can be fully regenerated by dehybridization, allowing repeated immobilization–measurement–regeneration cycles on the same chip across temperatures.

**Fig 4.**
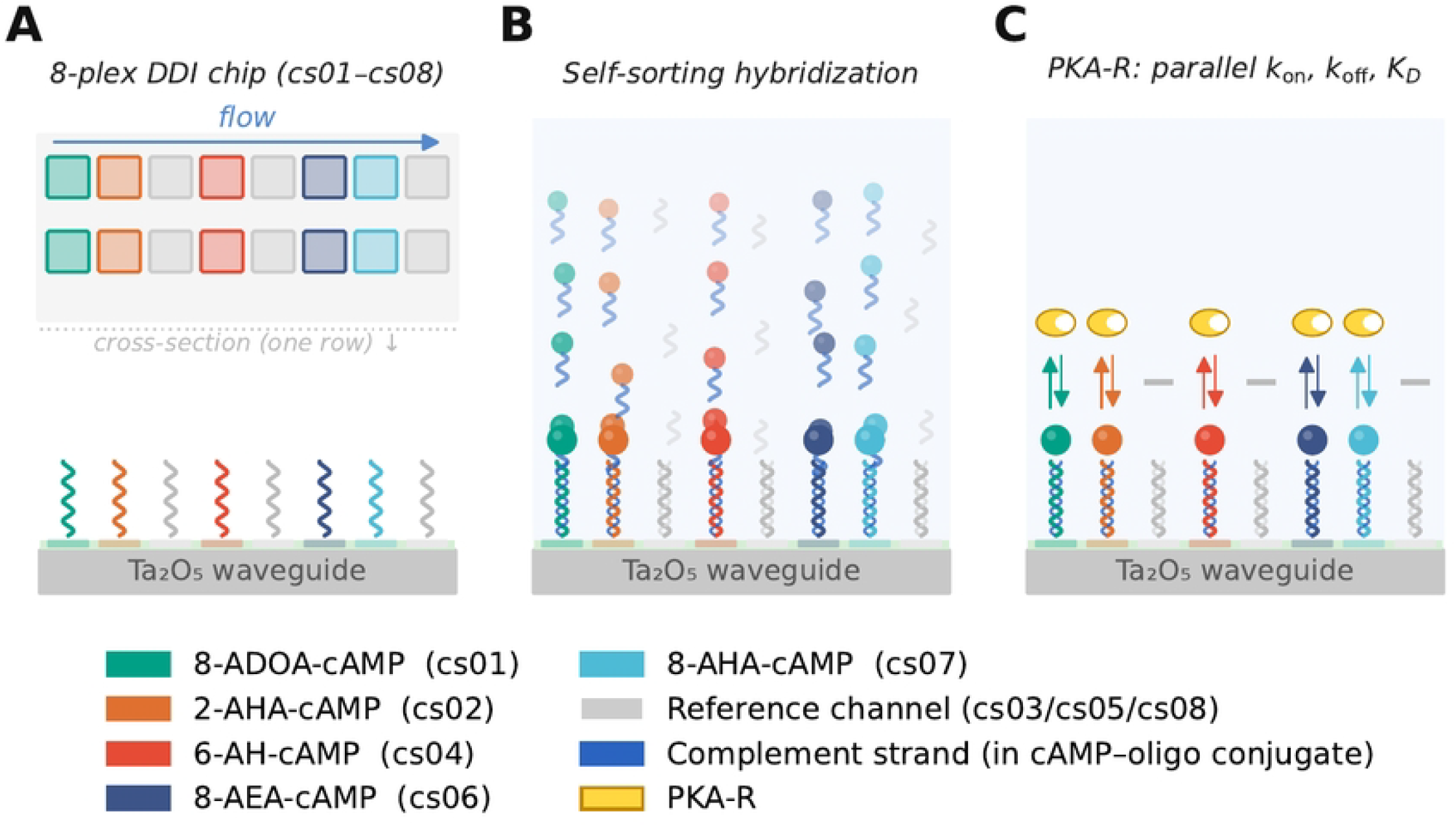
DNA-directed immobilization (DDI) multiplexing concept on an 8-plex focal molography chip. **(A)** Top view and cross-section of the 8-plex chip, showing sixteen molograms arranged in two rows of eight, with two molograms per capture-strand position (cs01–cs08), on a Ta_2_O_5_ waveguide coated with a PAA-*g* –PEG non-fouling brush. Five molograms carry sequence-specific capture strands for active analyte binding (cs01, cs02, cs04, cs06, cs07); three serve as negative controls (cs03, cs05, cs08). **(B)** A single injection step (200 nM each) containing all five cAMP–oligonucleotide conjugates and three reference oligonucleotides simultaneously. Each conjugate self-sorts by Watson–Crick hybridization to its complementary capture strand, loading the five active molograms with different cAMP derivatives in a single step. **(C)** Injection of PKA regulatory subunit (PKA-R) over the functionalized chip. PKA-R binds selectively to the immobilized cAMP derivatives; association (*k*_on_) and dissociation (*k*_off_) are measured simultaneously across all five compounds in one experiment, while negative controls carry non-complementary capture strands and report any non-specific binding to the ds-DNA backbone.

### Comparative thermodynamic fingerprinting of five cAMP derivatives by DDI-SCK

DDI experiments were performed in buffer A using the same 8-plex chip format; within each run, all five cAMP derivatives were carried simultaneously on the same chip. Five capture-strand positions carried sequence-specific capture strands for 6-AH-cAMP, 8-AHA-cAMP, 2-AHA-cAMP, 8-AEA-cAMP, and 8-ADOA-cAMP, while three positions served as negative controls with non-complementary capture strands (Fig 4A). A single injection containing all five cAMP–oligonucleotide conjugates loaded each chip by self-sorting hybridization (Fig 4B), achieving immobilization levels of 20–45 pg mm^*−*2^ depending on the derivative (S4 Table), after which PKA-R was injected and binding to all five derivatives was measured simultaneously (Fig 4C). Across the five temperatures (15–35°C), repeated DDI-SCK cycles were retained after quality filtering, giving *N* = 10, 18, 14, 10 and 12 individual fits per compound at 15, 20, 25, 30 and 35°C, respectively (Fig 5A, B; see Methods). 6-AH-cAMP, 8-AHA-cAMP, and 8-AEA-cAMP show well-behaved 1:1 Langmuir kinetics across all temperatures (Fig 5C, D, G), while 8-ADOA-cAMP and 2-AHA-cAMP exhibit a non-dissociating fraction and were fitted with a Langmuir 1:1 NonDissRelative model to account for this sticking component (Fig 5E, F; see Methods). Kinetic parameters (*k*_on_, *k*_off_, *K*_D_) at 25 °C for all five derivatives are summarized in Table 5. The negative controls show no detectable PKA-R binding (S4 Fig (c)), supporting the specificity of the DDI approach. Complete raw CMD sensorgrams and per-sensor Langmuir fits for every DDI-SCK run across all five temperatures are compiled in S7 Fig (a)–(af).

**Table 5.** Kinetic parameters for PKA-R binding to five cAMP derivatives at 25 °C measured by DDI-SCK.

| Ligand | $k_{\text{on}} (\times 10^5 \text{ M}^{-1} \text{ s}^{-1})$ | $k_{\text{off}} (\times 10^{-3} \text{ s}^{-1})$ | $K_D$ (nM) | $N$ |
| --- | --- | --- | --- | --- |
| 6-AH-cAMP | $7.67 \pm 0.85$ | $7.57 \pm 0.82$ | $10.0 \pm 1.4$ | 14 |
| 8-AHA-cAMP | $4.66 \pm 0.61$ | $4.32 \pm 0.42$ | $9.4 \pm 1.5$ | 14 |
| 8-ADOA-cAMP | $8.39 \pm 0.86$ | $13.74 \pm 1.67$ | $16.6 \pm 2.8$ | 14 |
| 2-AHA-cAMP | $2.98 \pm 0.46$ | $43.86 \pm 5.07$ | $150.3 \pm 29.2$ | 14 |
| 8-AEA-cAMP | $5.95 \pm 0.54$ | $5.65 \pm 0.68$ | $9.6 \pm 1.6$ | 14 |

Values are arithmetic mean ± sample standard deviation from the 14 retained individual DDI-SCK fits at 25 °C (seven included SCK cycles, two positive molograms per compound per cycle). $K_D = k_{\text{off}}/k_{\text{on}}$ . 8-ADOA-cAMP and 2-AHA-cAMP were fitted with a Langmuir 1:1 NonDissRelative model to account for a non-dissociating fraction; the $k_{\text{on}}$ , $k_{\text{off}}$ , and $K_D$ values reported here refer to the dissociating fraction only.

**Fig 5.**
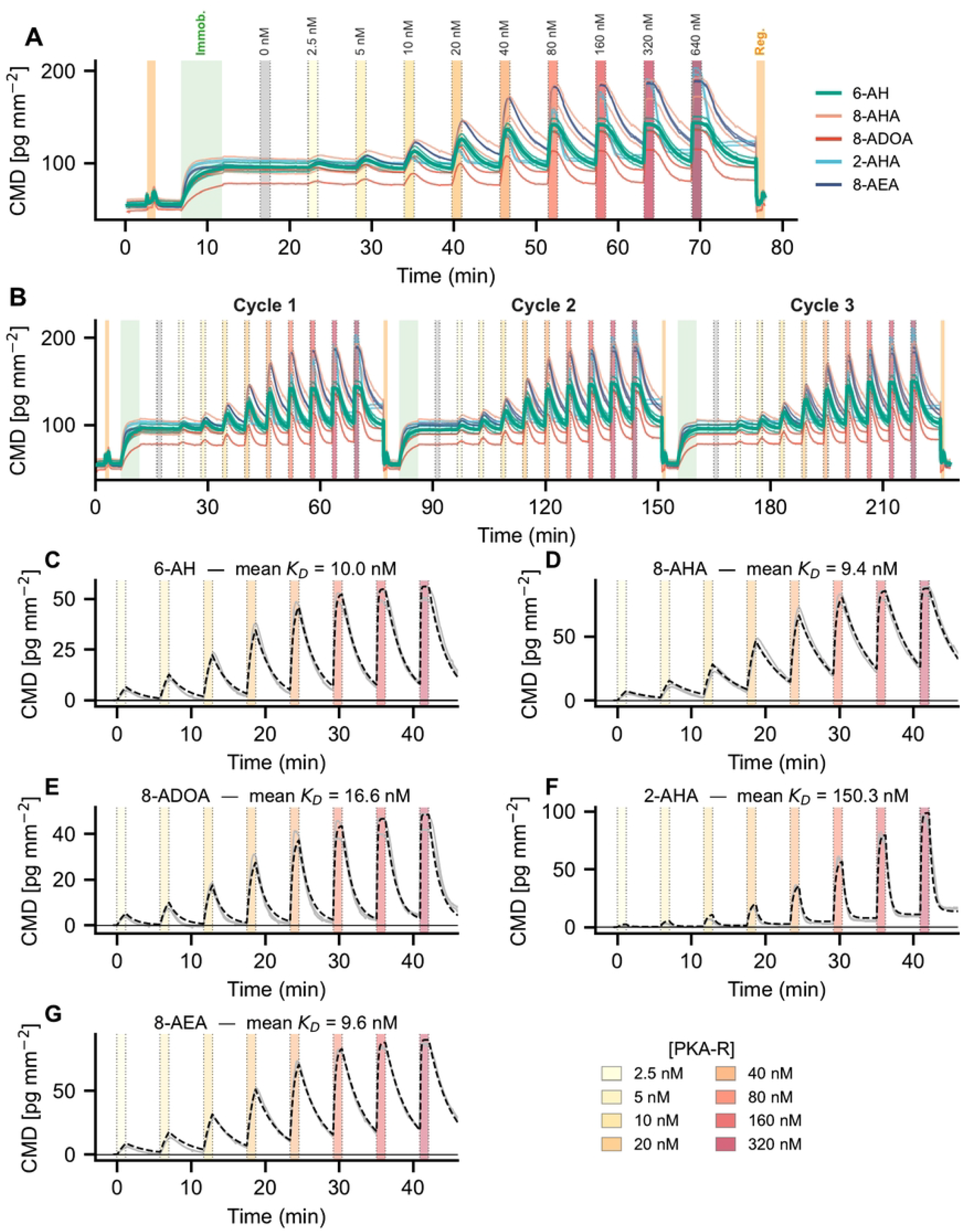
Representative DNA-directed immobilization single-cycle kinetics (DDI-SCK) experiment and example kinetic fits in buffer A. **(A)** Single complete DDI-SCK cycle at 25 °C showing DNA-directed capture of 6-AH-cAMP on an 8-plex chip, followed by ten ascending PKA-R concentration steps (0–640 nM) and a final regeneration step with 3 M GuHCl and 125 mM NaOH. The CMD signal (pg/mm^2^) is shown for both active sensors. Band labels denote the immobilization (Immob., green) and regeneration (Reg., orange) phases. **(B)** Full three-cycle DDI-SCK experiment at 25 °C illustrating reproducibility across three independent immobilization– measurement–regeneration cycles. **(C–G)** Representative kinetic fits for all five cAMP derivatives at 25 °C: **(C)** 6-AH-cAMP, **(D)** 8-AHA-cAMP, **(E)** 8-ADOA-cAMP, **(F)** 2-AHA-cAMP, **(G)** 8-AEA-cAMP. Traces show CMD offset to the pre-injection baseline immediately before each concentration step (display offset only; not blank-cycle subtracted). Individual sensor traces (gray) are overlaid with the mean 1:1 Langmuir fit (black dashed). Pink shading indicates PKA association steps. The apparent equilibrium dissociation constant *K*_*D*_ at 25 °C is annotated.

Isoaffinity plots at each temperature (Fig 6A–E) reveal that four of the five compounds — 6-AH-cAMP, 8-AHA-cAMP, 8-AEA-cAMP, and 8-ADOA-cAMP — cluster in a similar *K*_D_ range (~10–20 nM), but achieve this affinity through distinct combinations of *k*_on_ and *k*_off_. At some temperatures (notably 15 and 30 °C), distinct sub-clusters are visible within the individual-fit point clouds, most likely reflecting run-to-run variation in effective analyte concentration arising from pipetting imprecision or partial degradation of the protein stock between DDI-SCK cycles; this intra-temperature scatter is incorporated into the weighted analysis and contributes to the fitted uncertainty. The exception is 2-AHA-cAMP, which exhibits substantially weaker affinity (*K*_D_ ≈150–300 nM; Δ*G* = −9.2 kcal/mol vs. −10.5 to −10.9 kcal/mol for the others; Table 6), consistent with modification at the C2 position of the adenine ring affecting the binding pocket interaction differently than modification at the N^6^ or C8 positions.

**Table 6.** Summary of equilibrium thermodynamic parameters for DNA-directed immobilization (DDI-SCK).

| Ligand | Format | Matrix | Dataset | $\Delta H$<br>(kcal/mol) | $-T\Delta S$<br>(kcal/mol) | $\Delta G$<br>(kcal/mol) | $K_D$<br>(nM) | $R^2$ | $N$ |
| --- | --- | --- | --- | --- | --- | --- | --- | --- | --- |
| 6-AH-cAMP | On-DNA | Buffer A | 15–35°C | $-5.33 \pm 0.75$ | $-5.55 \pm 0.75$ | $-10.88 \pm 0.02$ | $10.6 \pm 0.4$ | 0.635 | 64 |
| 8-AHA-cAMP | On-DNA | Buffer A | 15–35°C | $-5.11 \pm 0.89$ | $-5.79 \pm 0.89$ | $-10.90 \pm 0.02$ | $10.2 \pm 0.3$ | 0.780 | 64 |
| 8-ADOA-cAMP | On-DNA | Buffer A | 15–35°C | $-7.39 \pm 0.84$ | $-3.14 \pm 0.85$ | $-10.54 \pm 0.02$ | $18.8 \pm 0.6$ | 0.730 | 64 |
| 2-AHA-cAMP | On-DNA | Buffer A | 15–35°C | $-7.18 \pm 1.24$ | $-2.03 \pm 1.25$ | $-9.21 \pm 0.03$ | $177 \pm 9$ | 0.663 | 64 |
| 8-AEA-cAMP | On-DNA | Buffer A | 15–35°C | $-6.96 \pm 0.99$ | $-3.96 \pm 0.98$ | $-10.91 \pm 0.02$ | $10.1 \pm 0.3$ | 0.898 | 64 |

Weighted Van’t Hoff fits across 5 temperatures (15–35°C). Equilibrium parameters use the 1 M standard-state convention ( $K_D/C^\circ$ , $C^\circ = 1$ M). Central values were obtained from linear fits to $\chi^2$ -weighted per-temperature means; uncertainties are bootstrap standard deviations from resampling retained fits within each temperature group. $\Delta H$ and $\Delta S$ are extracted from the Van’t Hoff fit (temperature-independent parameters); $\Delta G = \Delta H - T\Delta S$ is evaluated at $T_{\text{ref}} = 25^\circ\text{C}$ (298.15 K). $K_D$ is the dissociation constant at $T_{\text{ref}} = 25^\circ\text{C}$ , obtained from $\Delta G$ as $K_D = C^\circ \exp(\Delta G/RT)$ ( $C^\circ = 1$ M); its uncertainty is propagated from $\Delta G$ . $N$ = total number of retained individual DDI-SCK fits across all temperatures (15°C: 10, 20°C: 18, 25°C: 14, 30°C: 10, 35°C: 12; total 64 per compound).

**Fig 6.**
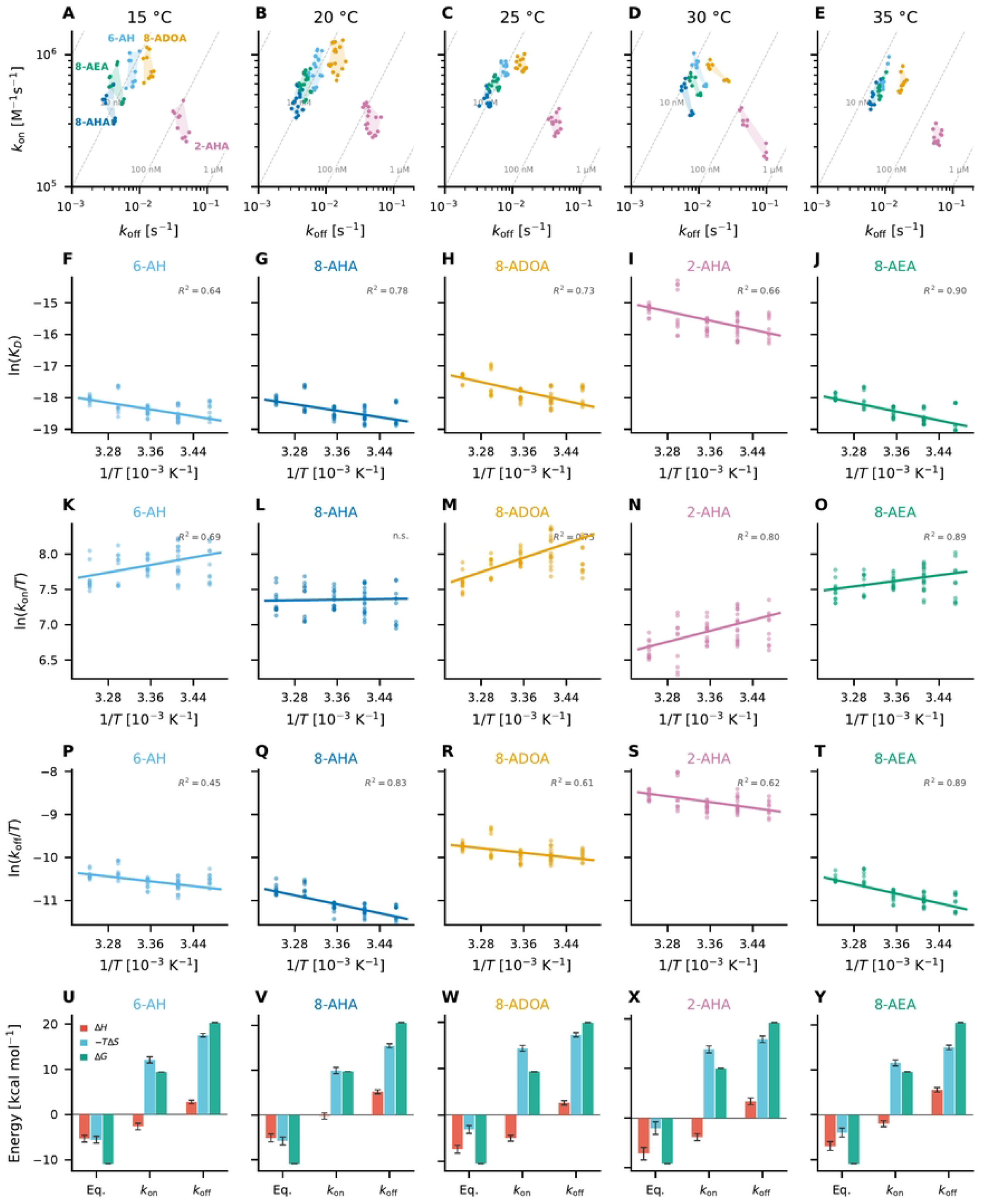
Comparative thermodynamic profiling of five cAMP derivatives using DNA-directed immobilization single-cycle kinetics (DDI-SCK) in buffer A. **(A–E)** Isoaffinity scatter plots of *k*_off_ vs. *k*_on_ for all five cAMP derivatives at 15, 20, 25, 30, and 35 °C, respectively. Diagonal dashed lines indicate isoaffinity contours for constant *K*_*D*_ values. Shaded regions represent the convex hull of individual retained DDI-SCK fits for each compound. Compound labels are shown in panel A. **(F–J)** Weighted Van’t Hoff analysis using a 1 M standard state (ln(*K*_*D*_*/C*°) vs. 1*/T*, with *C*° = 1 M) per compound. Individual retained fit results are shown as points; the line and annotated parameters are from the *χ*^2^-weighted five-temperature analysis described in Methods. The coefficient of determination (*R*^2^) is indicated. **(K–O)** Eyring analysis of the association rate constant using a 1 M standard state (ln(*k*_on_*C*°*/T*) vs. 1*/T*, with *C*° = 1 M). **(P–T)** Eyring analysis of the dissociation rate constant (ln(*k*_off_ */T*) vs. 1*/T*). **(U–Y)** Thermodynamic decomposition of binding free energy (Δ*G*), enthalpy (Δ*H*), and entropic contribution (*−T* Δ*S*) from equilibrium Van’t Hoff (Eq.), Eyring *k*_on_, and Eyring *k*_off_ analyses. Error bars represent bootstrap standard deviations from resampling retained fits within each temperature group.

Van ‘t Hoff analysis yields distinct thermodynamic fingerprints for each derivative (Fig 6F–J; Table 6): all five show exothermic binding (Δ*H*° ranging from −5.1 to −7.4 kcal/mol), but the balance between enthalpic and entropic contributions differs. 6-AH-cAMP and 8-AHA-cAMP show a roughly equal split between Δ*H* and −*T* Δ*S*, whereas 2-AHA-cAMP is more strongly enthalpy-driven with a smaller entropic contribution. 8-AEA-cAMP and 8-ADOA-cAMP fall between these extremes. The Eyring analyses of *k*_on_ and *k*_off_ (Fig 6K–T; Table 7) provide the activation-level decomposition and are largely consistent with the equilibrium picture. One exception is the association Eyring fit for 8-AHA-cAMP, whose slope is statistically indistinguishable from zero: the derived activation enthalpy 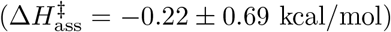 is consistent with no temperature dependence (0.3 *σ*), so the plot of ln(*k*_on_*C*°*/T*) versus 1*/T* is essentially flat and the weighted *R*^2^ (0.006), which is ill-defined for a zero-slope fit, falls marginally below zero. Association of this derivative therefore has no resolvable enthalpic activation barrier across the 15–35°C range, and 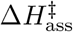 for 8-AHA-cAMP should not be interpreted quantitatively. The activation free energy 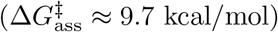 remains well-determined from the intercept and is almost entirely entropic 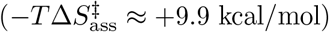, reflecting the entropic cost of forming the encounter complex under the 1 M association standard state. The thermodynamic decomposition bar charts (Fig 6U–Y) summarize the energetic fingerprints derived from all three analyses and confirm that each derivative exhibits a distinct but internally consistent thermodynamic profile.

**Table 7.**
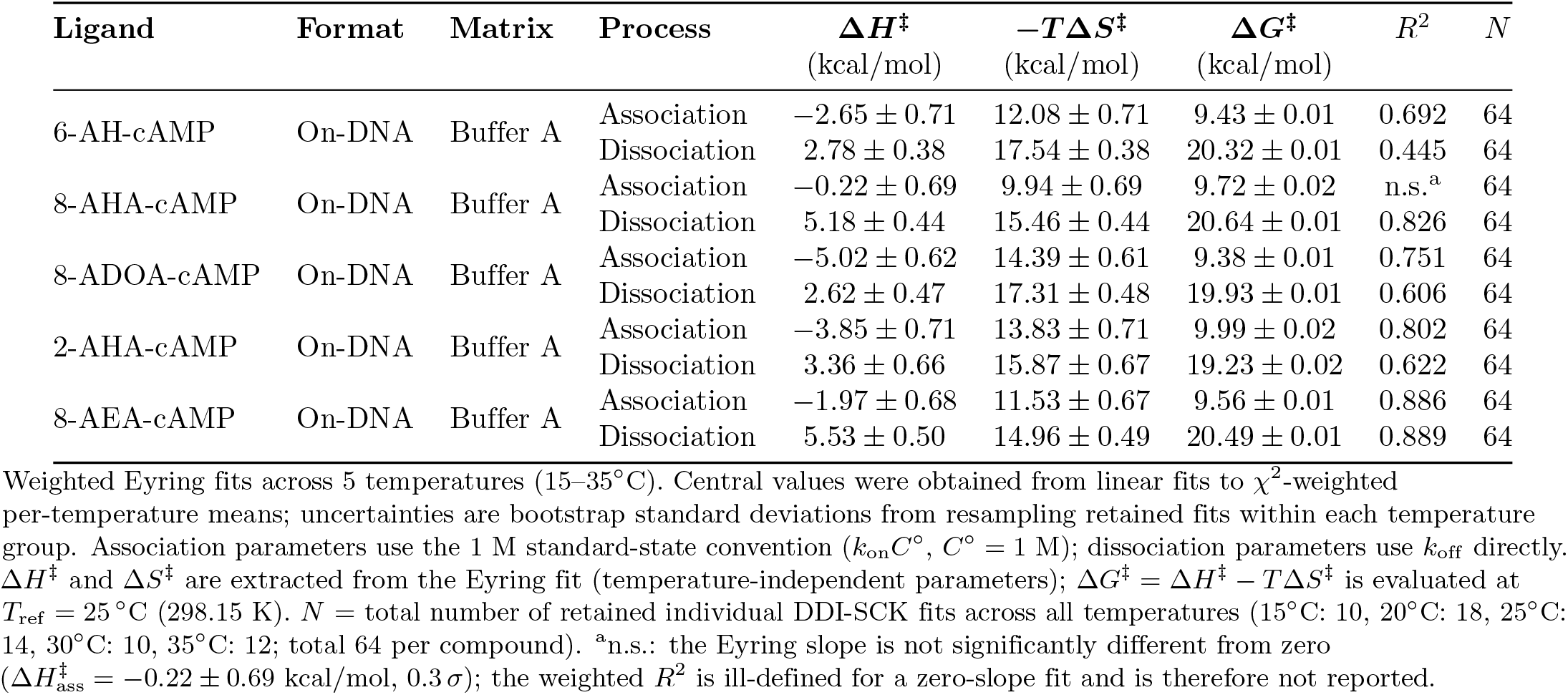
Summary of activation parameters from Eyring analysis.

These results demonstrate that DDI on focal molography enables comparative thermodynamic fingerprinting — multiple ligands profiled side by side within the same multiplexed run under closely matched conditions — reducing chip-to-chip variability as a confounding factor in the comparative analysis.

## Discussion

### What the study demonstrates

Focal molography was originally developed as a label-free *kinetic* biosensor [9, 27]. Here we show that the same kinetic readout can be used for apparent thermodynamic characterization: by acquiring single-cycle binding kinetics across a range of temperatures and applying van ‘t Hoff and Eyring analyses, focal molography yields apparent equilibrium (Δ*G*, Δ*H*, −*T* Δ*S*) and activation (Δ*G*^*‡*^, Δ*H*^*‡*^, *T* −Δ*S*^*‡*^) parameters for a protein–ligand interaction. Combined with DNA-directed immobilization (DDI), which displays several ligands on a multiplexed chip, the approach lets these thermodynamic fingerprints be compared across multiple compounds measured side by side within the same experimental framework. We stress that this is a demonstration of a measurement *workflow*, not a claim that focal molography reproduces absolute solution thermodynamics: the parameters reported here are apparent, format-specific quantities, most powerful for ranking and contrasting related ligands rather than as substitutes for calorimetric reference values. The property that makes the workflow practical—and distinguishes it from refractometric surface-based methods—is the intrinsic temperature insensitivity of the diffractometric readout, which we consider next.

### Why focal molography is suited to temperature-dependent and matrix-tolerant kinetic measurements

The diffractometric (coherent mass density, CMD) channel encodes binding in a lithographically defined grating and reads it out against the sub-micron self-reference of that pattern, strongly suppressing sensitivity to the bulk refractive-index changes that dominate refractometric surface readouts [18, 19]. During a continuous 25–45–25 °C temperature ramp, the CMD baseline drifts by less than 1 pg mm^*−*2^, whereas the refractometric (mass density, MD) channel—even with active sensor-pair referencing (MD_ref_)—shows residual excursions of up to ~±1200 pg mm^*−*2^ (Fig 2A–C). The diffractometric readout therefore avoids the lengthy per-step thermal equilibration that limits temperature ramping on refractometric platforms [2], making multi-temperature kinetic acquisition practical within a single session. The same self-referencing also suppresses bulk refractive-index effects and much of the spatially incoherent non-specific background: in 50% human serum the raw CMD sensorgrams resolve the concentration-dependent binding signature (Fig 2D, E), despite measurable non-specific CMD adsorption of ~ 16 pg mm^*−*2^ from this serum preparation in blank controls (S4 Fig (b)B), whereas the refractometric channel is overwhelmed by serum signals that persist even after blank subtraction (Fig 2F, G). These properties suppress a major class of bulk-refractive-index and non-specific artifacts in the CMD channel rather than eliminating all sources of error.

The 20 °C covalent-format MCK experiments further show that this analytical robustness translates into quantitative kinetic measurements in a complex matrix. For both 6-AH-cAMP and 8-AHA-cAMP, the association rates measured in 50% human serum were essentially unchanged relative to buffer A, while *k*_off_ and *K*_*D*_ were only modestly higher, by approximately 20% (Table 4). Thus, under the conditions tested, serum did not obscure the specific PKA-R–cAMP interaction or produce a qualitatively different kinetic signature. This result should be interpreted with appropriate scope: the serum comparison was performed at a single temperature, in the covalent immobilization format, and with two derivatives only. It supports matrix-tolerant kinetic analysis by the CMD channel, whereas the temperature-dependent DDI thermodynamic profiling reported here was performed in buffer A. The contrasting behavior of the TSH-depleted serum used on the prototype instrument and the normal human serum used on the MACS^®^ Matchmaker further indicates that non-specific adsorption remains serum-lot- and surface-dependent and should be assessed for each experimental context.

### What DNA-directed immobilization adds

DNA-directed immobilization is what makes the comparative profiling practical. Its central benefit is that, within each DDI-SCK run, all five conjugates are loaded onto the same 8-plex chip by self-sorting hybridization and measured under closely matched conditions—shared chip format, analyte injection, and temperature history within that run—so that differences in Δ*H* and −*T*Δ*S* between compounds are substantially less confounded by chip-to-chip and run-timing variability than separate single-ligand experiments would be (Fig 6). The hybridized layer is also regenerable: stripping the conjugates and bound analyte lets the same chip be reloaded and re-measured in repeated immobilize–measure–regenerate cycles at each temperature. Combined with the single-cycle format, which removes regeneration between concentration steps, this makes the workflow fast: once the cAMP–oligonucleotide conjugates are prepared, a complete five-temperature profile for five compounds can be acquired within a single day per chip/run. The self-sorting step was robust throughout, with a single 5-minute injection loading all five conjugates to functional levels (S4 Table), while three non-complementary control spots showed no detectable non-specific PKA-R binding to the DNA-presenting control surfaces (S4 Fig (c)).

These gains are not free. DDI does not remove all measurement variability—residual run-to-run scatter, for example in effective analyte concentration, persists and is incorporated into the weighted fitting and the resulting uncertainty—and it adds preparation overhead, since each ligand must be conjugated to an oligonucleotide by click chemistry and purified, roughly one to two days of bench work beyond direct covalent coupling. For routine kinetic screening, where the thermodynamic decomposition is not the readout, covalent immobilization remains simpler and sufficient. Where the enthalpic and entropic contributions are themselves the quantities of interest—as in thermodynamic fingerprinting for lead optimization or mechanistic work—the controlled, regenerable, multiplexed presentation of DDI can justify the extra effort. We therefore recommend DDI as the default architecture for comparative thermodynamic profiling by focal molography, while treating the absolute thermodynamic values it yields as specific to the DNA-tethered format.

### Apparent thermodynamic fingerprints of the five cAMP derivatives

Interpreted as apparent, format-specific parameters, the five derivatives nonetheless show resolvable, distinct apparent thermodynamic patterns (Table 6). Four of them—6-AH-, 8-AHA-, 8-AEA-, and 8-ADOA-cAMP—bind with closely similar affinity (Δ*G* between −10.5 and −10.9 kcal/mol), while 2-AHA-cAMP is a clear outlier, roughly 1.5 kcal/mol weaker (Δ*G* = −9.2 kcal/mol). This is consistent with the chemistry of linker attachment: 2-AHA-cAMP is modified at the C2 position of the adenine ring, whereas the others carry their linkers at the N^6^ or C8 positions, and C2 substitution would be expected to perturb the interaction differently—though we treat this as a correlation with linker position rather than a structurally resolved mechanism. Notably, the four high-affinity derivatives do not reach their similar Δ*G* by the same route: binding is exothermic throughout (Δ*H* from 5.1 to −7.4 kcal/mol) and accompanied by a favorable entropy term, expressed here as negative −*T* Δ*S*. The balance ranges from near-equal enthalpic and entropic contributions for 6-AH- and 8-AHA-cAMP (Δ*H* ≈ −*T* Δ*S* ≈ −5 kcal/mol) to a more enthalpy-weighted profile for 8-AEA- and 8-ADOA-cAMP. The weaker-binding 2-AHA-cAMP shows the smallest favorable entropy term (−*T* Δ*S* = −2.0 kcal/mol) and therefore the strongest relative enthalpic weighting, even though its absolute Δ*H* is similar to that of 8-ADOA-cAMP. Affinity alone would have hidden these differences, which is precisely the information thermodynamic fingerprinting adds.

The Eyring decomposition of the rate constants is internally consistent with this picture (Table 7). For all derivatives, the association barrier is dominated by an unfavorable entropic contribution: 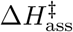 is small or negative (*−*0.2 to *−*5.0 kcal/mol), whereas 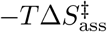 contributes +10 to +14 kcal/mol. In contrast, dissociation includes a positive enthalpic component 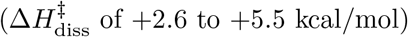. The one exception is the association fit for 8-AHA-cAMP, whose slope is statistically indistinguishable from zero, so its 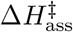 should not be read quantitatively (see below). Across the panel, these activation fingerprints reinforce the equilibrium observation: chemically similar ligands of comparable affinity can follow measurably different kinetic and energetic routes to the bound state.

### Interpretation boundaries and measurement limitations

Several boundaries constrain how these parameters should be read. First, the thermodynamic profiling was carried out in buffer A only; the serum measurements were limited to single-temperature kinetics in the covalent format, so we make no claim about temperature-dependent thermodynamics in complex matrices. Second, the DDI architecture itself is part of the measured system: the DNA tether changes ligand distance from the surface, local charge and hydration, and the accessible orientational ensemble, so the resulting rate and thermodynamic parameters are apparent parameters of the DNA-tethered presentation. Third, the measurements underlying each thermodynamic analysis (*N* = 64 per compound) are technical and assay replicates—multiple sensors, cycles, and experimental sessions on the same chip format and protein preparation—rather than fully independent biochemical replicates, so the bootstrap uncertainties mainly capture within-assay measurement and surface variability, not between-preparation reproducibility. Fourth, 8-ADOA-cAMP and 2-AHA-cAMP displayed a non-dissociating fraction and were fitted with a Langmuir 1:1 NonDissRelative model; their reported rate and thermodynamic parameters describe the dissociating population only and are correspondingly model-dependent. Fifth, visible intra-temperature dispersion remains in the data (Fig 6A–E), plausibly including run-to-run differences in effective analyte concentration; this scatter contributes to the weighted van ‘t Hoff and Eyring fits and to the bootstrap uncertainty, but is not removed by the analysis. Finally, the van ‘t Hoff and Eyring analyses assume temperature-independent equilibrium and activation enthalpies and entropies across the 15–35 °C window; any curvature, including heat-capacity effects, would be folded into the linear-fit parameters. For all these reasons, the values reported here are best treated as apparent, format-specific thermodynamic fingerprints for comparison across ligands, not as absolute thermodynamic constants for the cAMP–PKA-R interaction.

One fit warrants a specific caveat. For 8-AHA-cAMP the association Eyring slope is not significantly different from zero 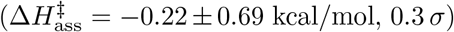, which leaves the weighted *R*^2^ ill-defined for that fit; the activation enthalpy of association for this derivative therefore carries no resolvable temperature dependence and should not be interpreted quantitatively. The corresponding activation free energy 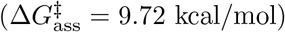 remains well constrained because it is determined by the fitted association rate near the reference temperature; however, the enthalpy–entropy split for this association process should be interpreted only qualitatively.

### Comparison with SPR and ITC

To place the apparent FM-DDI fingerprints in context, we compared the two derivatives for which published SPR and ITC data are available, 6-AH-cAMP and 8-AHA-cAMP, with the measurements of Moll et al. [21] (Fig 7; Tables 8 and 9). This comparison should be read as external context rather than as a test of thermodynamic equivalence: the three methods differ in ligand presentation, surface chemistry, analysis workflow, and, for ITC, the absence of immobilization. Within that scope, the equilibrium free energies fall in the expected nanomolar-binding range. FM-DDI, SPR, and ITC agree in Δ*G* to within approximately 1–1.5 kcal/mol, consistent with the kinetic fits capturing the same specific PKA-R–cAMP recognition event. At the same time, this level of agreement should not be over-interpreted, because a 1.4 kcal/mol shift corresponds to roughly an order-of-magnitude change in *K*_*D*_ at room temperature. The absolute affinities therefore still differ appreciably between platforms, most clearly for 8-AHA-cAMP, where the published SPR value (110 nM) is much weaker than the direct 25 °C FM-DDI value (9.4 nM).

**Fig 7.**
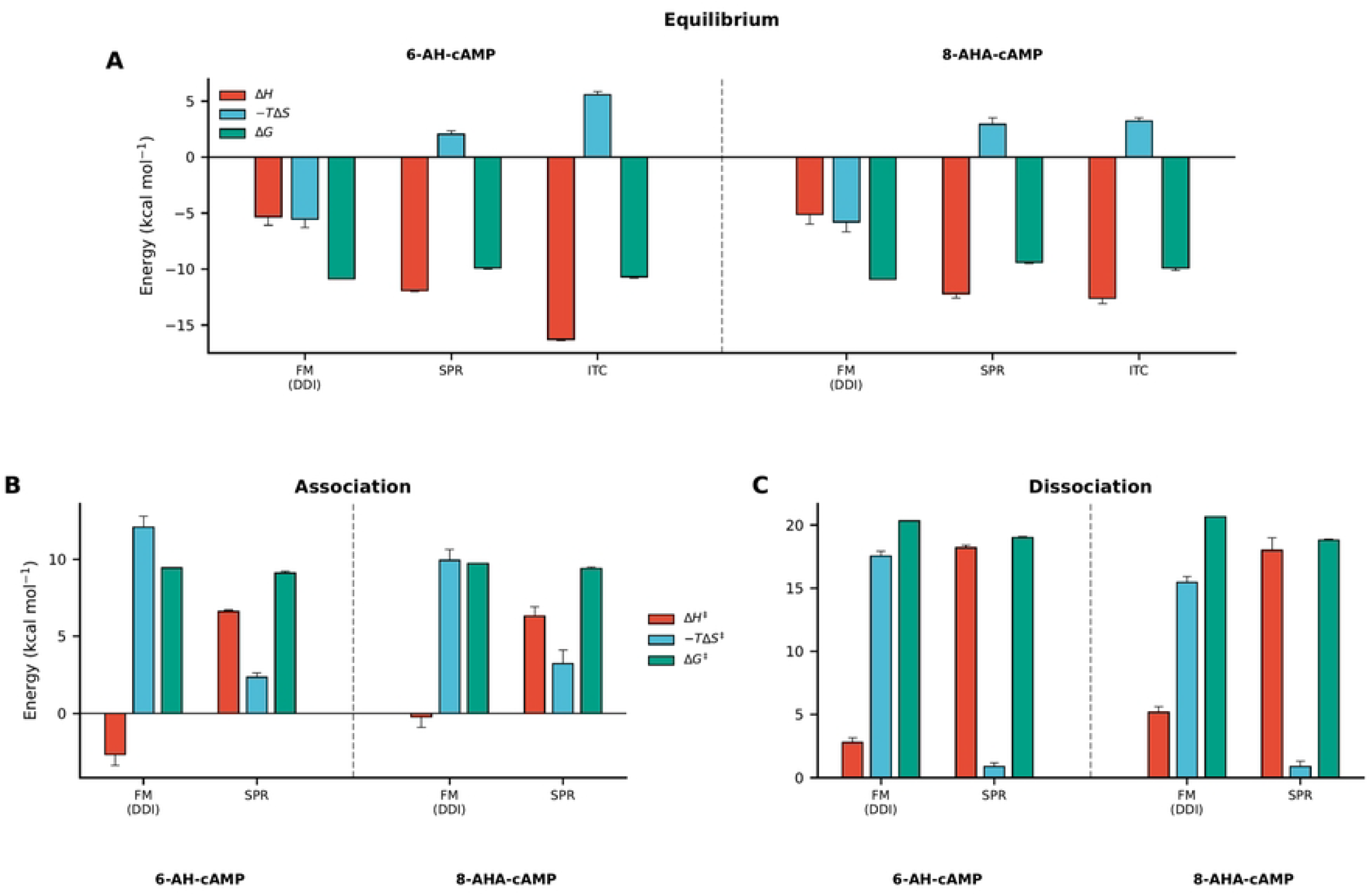
Thermodynamic and activation parameter comparison across measurement techniques. **(A)** Equilibrium thermodynamic decomposition: binding free energy (Δ*G*, teal), enthalpy (Δ*H*, red), and entropic contribution (−*T* Δ*S*, blue) for **6-AH-cAMP** (left group) and **8-AHA-cAMP** (right group) measured by focal molography with DNA-directed immobilization (FM DDI), SPR, and ITC. Δ*G* values are broadly consistent across methods (within ~1–1.5 kcal/mol), whereas Δ*H* and −*T* Δ*S* show method-dependent decompositions: FM DDI is more entropy-balanced, while the SPR and ITC values are more strongly enthalpy-driven. **(B)** Activation parameters for **association**: activation free energy (Δ*G*^*‡*^, teal), activation enthalpy (Δ*H*^*‡*^, red), and −*T* Δ*S*^*‡*^ (blue). FM and SPR yield comparable Δ*G*^*‡*^, while the enthalpic/entropic decomposition differs. **(C)** Activation parameters for **dissociation**: the most striking cross-platform difference appears here, where SPR reports a much larger Δ*H*^*‡*^ and correspondingly smaller −*T* Δ*S*^*‡*^ than focal molography, consistent with method-dependent differences in ligand presentation and local surface environment. Error bars for FM DDI represent bootstrap standard deviations from this study; external-method error bars represent replicate uncertainties reported by Moll et al. [21]: SPR and ITC in panel A, and SPR only in panels B and C.

**Table 8.** Equilibrium thermodynamic parameters at 298 K and representative direct *K*_*D*_ values for PKA-R binding to 6-AH-cAMP and 8-AHA-cAMP across measurement techniques. All values are buffer-phase (no serum): FM DDI was measured in buffer A (this study), while the SPR and ITC values are from Moll et al. in their respective study buffers.

| Ligand | Method | $K_D$ (nM) | $\Delta G$ | $\Delta H$ | $-T\Delta S$ |
| --- | --- | --- | --- | --- | --- |
|  |  |  | (kcal/mol) | (kcal/mol) | (kcal/mol) |
| 6-AH-cAMP | FM DDI | $10.0 \pm 1.4^a$ | $-10.88 \pm 0.02$ | $-5.33 \pm 0.75$ | $-5.55 \pm 0.75$ |
| | SPR | $38 \pm 5$ | $-9.90 \pm 0.1$ | $-11.9 \pm 0.1$ | $+2.05 \pm 0.29$ |
| | ITC | $10 \pm 1$ | $-10.70 \pm 0.1$ | $-16.3 \pm 0.1$ | $+5.57 \pm 0.29$ |
| 8-AHA-cAMP | FM DDI | $9.4 \pm 1.5^a$ | $-10.90 \pm 0.02$ | $-5.11 \pm 0.89$ | $-5.79 \pm 0.89$ |
| | SPR | $110 \pm 21$ | $-9.40 \pm 0.1$ | $-12.2 \pm 0.4$ | $+2.93 \pm 0.59$ |
| | ITC | $46 \pm 15$ | $-9.90 \pm 0.2$ | $-12.6 \pm 0.5$ | $+3.22 \pm 0.29$ |

FM = focal molography; DDI = DNA-directed immobilization. FM DDI thermodynamic values from this study (Table 6); FM DDI thermodynamic uncertainties are bootstrap standard deviations as defined in Methods. SPR and ITC values from Moll et al. [21]. <sup>a</sup>FM $K_D$ values are direct Langmuir 1:1 kinetic-fit means at $T = 25^\circ\text{C}$ (298.15 K), reported as mean ± standard deviation across 14 retained individual DDI-SCK fits; they are not back-calculated from $\Delta G$ . FM $\Delta G$ , $\Delta H$ , and $-T\Delta S$ are evaluated from Van't Hoff analysis at $T_{\text{ref}} = 25^\circ\text{C}$ (298.15 K); SPR and ITC thermodynamic parameters from Moll et al. are also reported at 298 K. $\Delta G$ values are broadly consistent across methods (agreeing to within $\sim 1\text{--}1.5$ kcal/mol), but the $\Delta H/-T\Delta S$ decomposition differs by method: FM DDI is more entropy-balanced, whereas the SPR and ITC values are more strongly enthalpy-driven.

**Table 9.**
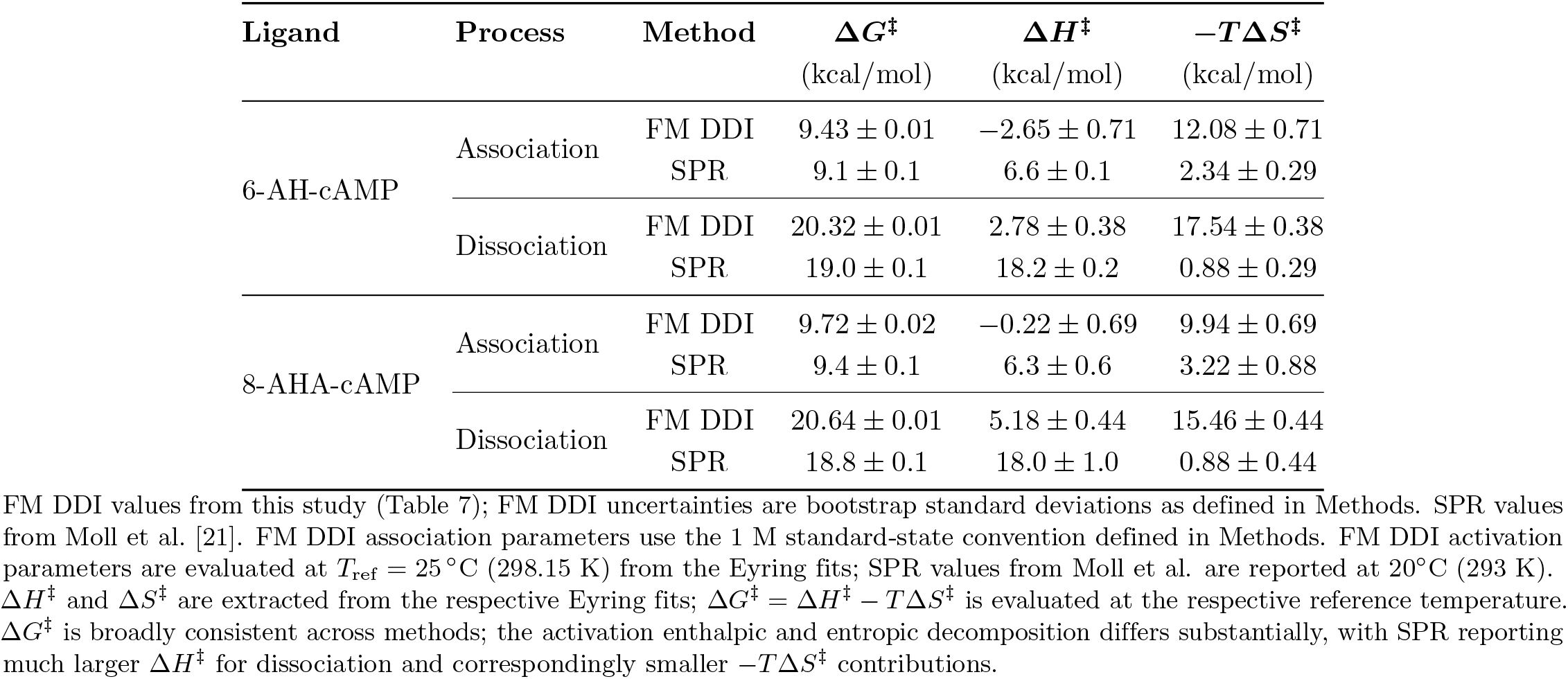
Activation parameters (Δ*G*^*‡*^, Δ*H*^*‡*^, *−T* Δ*S*^*‡*^) for association and dissociation of PKA-R with AH-cAMP and 8-AHA-cAMP across measurement techniques (reference temperatures vary by method; see footnotes). All values are buffer-phase (no serum): FM DDI was measured in buffer A (this study), while the SPR values are from Moll et al. in their respective study buffer.

| Ligand | Process | Method | $\Delta G^\ddagger$<br>(kcal/mol) | $\Delta H^\ddagger$<br>(kcal/mol) | $-T\Delta S^\ddagger$<br>(kcal/mol) |
| --- | --- | --- | --- | --- | --- |
| 6-AH-cAMP | Association | FM DDI | $9.43 \pm 0.01$ | $-2.65 \pm 0.71$ | $12.08 \pm 0.71$ |
| | | SPR | $9.1 \pm 0.1$ | $6.6 \pm 0.1$ | $2.34 \pm 0.29$ |
| | Dissociation | FM DDI | $20.32 \pm 0.01$ | $2.78 \pm 0.38$ | $17.54 \pm 0.38$ |
| | | SPR | $19.0 \pm 0.1$ | $18.2 \pm 0.2$ | $0.88 \pm 0.29$ |
| 8-AHA-cAMP | Association | FM DDI | $9.72 \pm 0.02$ | $-0.22 \pm 0.69$ | $9.94 \pm 0.69$ |
| | | SPR | $9.4 \pm 0.1$ | $6.3 \pm 0.6$ | $3.22 \pm 0.88$ |
| | Dissociation | FM DDI | $20.64 \pm 0.01$ | $5.18 \pm 0.44$ | $15.46 \pm 0.44$ |
| | | SPR | $18.8 \pm 0.1$ | $18.0 \pm 1.0$ | $0.88 \pm 0.44$ |

The size of this cross-method spread is analyte-dependent, which is consistent with ligand-presentation effects. In the focal molography data, 8-AHA-cAMP is more sensitive to immobilization format than 6-AH-cAMP: compared at a matched temperature of 20 °C, its dissociation rate is about 3.3-fold slower in the DNA-tethered format (*k*_off_ ≈ 3.9 ×10^*−*3^ s^*−*1^) than in the covalent format (≈12.8 ×10^*−*3^ s^*−*1^; Table 4), whereas the corresponding difference for 6-AH-cAMP is only about 1.4-fold. The same qualitative pattern is seen in the external comparison, where 8-AHA-cAMP shows the largest SPR-to-FM-DDI affinity difference. Moreover, both covalent surface formats—focal molography in the covalent MCK experiment and SPR on carboxymethyl-dextran— report weaker apparent affinity for 8-AHA-cAMP than the DNA-tethered FM-DDI format. A parsimonious interpretation is that the C8-linked derivative is particularly sensitive to how the linker presents the cAMP moiety to PKA-R, and that the DNA tether may partly relieve constraints present in direct covalent presentation. This remains a hypothesis: the present data do not distinguish between linker distance, orientation, local electrostatics, hydration, or rebinding effects.

The enthalpy–entropy decomposition reveals a different grouping. FM-DDI reports near-balanced binding for both analogues (Δ*H*≈− *T* Δ*S* ≈−5 kcal/mol), whereas the published SPR and ITC values are both more strongly enthalpy-driven and opposed by an entropic penalty (Δ*H* from approximately *−*12 to *−*16 kcal/mol; −*T* Δ*S* from +2 to +6 kcal/mol). For 8-AHA-cAMP, the SPR and ITC decompositions are nearly identical despite one method being surface-based and the other solution-phase. The split is therefore not simply surface-versus-solution, nor simply kinetic van ‘t Hoff analysis versus calorimetry. Instead, the FM-DDI decomposition should be understood as an apparent fingerprint of the molographic measurement format, including DNA tethering, ligand presentation, and the sparse, hydrated PEG-brush surface. This interpretation is consistent with a format-dependent local surface environment, but the present data do not identify a specific molecular mechanism. A similar pattern appears in the activation parameters: Δ*G*^*‡*^ values are broadly comparable between FM-DDI and SPR, but the enthalpic and entropic components differ substantially, especially for dissociation, where SPR reports a much larger Δ*H*^*‡*^ and a much smaller *−T* Δ*S*^*‡*^ contribution. Taken together, the cross-method comparison supports the qualitative validity of FM-DDI thermodynamic fingerprinting while reinforcing that the absolute enthalpy–entropy split is the least transferable part of the comparison.

### Positioning and outlook

We therefore position focal molography as a complementary addition to the established biophysical toolkit rather than a replacement for SPR or ITC. For high-precision calorimetric thermodynamics of purified components in solution, ITC remains the reference method; for high-throughput kinetic and affinity screening on well-characterized surfaces, SPR is mature and widely accessible. What neither offers easily is comparative, temperature-resolved thermodynamic profiling at high throughput: van ‘t Hoff and Eyring analysis across multiple temperatures by SPR remains comparatively rare, largely because each temperature step demands lengthy equilibration and re-referencing. Focal molography with DDI fills this niche—multiplexed, low-sample-consumption, rapid temperature-dependent kinetic measurement—and is strongest where the comparative ranking of enthalpic and entropic fingerprints across related ligands, rather than the absolute calorimetric decomposition of any one ligand, is the goal.

Several directions follow naturally. The most immediate is to extend DDI-based temperature-dependent profiling from buffer into serum and other complex matrices, building on the matrix tolerance of the CMD channel demonstrated here at a single temperature. Beyond that, the approach invites application to broader target classes beyond small-molecule–protein interactions, to larger and more systematically varied ligand series for structure–activity studies, and to a controlled, side-by-side comparison of covalent and DNA-tethered presentation to characterize the format dependence identified above. Finally, validating selected DDI thermodynamic trends across independent chips and comparing selected molographic fingerprints against orthogonal solution-phase measurements would help establish which features of the apparent thermodynamic signatures are transferable across methods.

## Conclusion

This work establishes focal molography as a platform for apparent thermodynamic profiling of biomolecular interactions from temperature-dependent kinetic measurements. By exploiting the intrinsic temperature stability of the diffractometric readout and its ability to resolve binding kinetics in 50% human serum at 20°C, we demonstrate a temperature- and matrix-tolerant kinetic readout for cAMP–PKA-R binding.

Combined with DNA-directed immobilization, focal molography enables multiplexed comparative profiling: using the same 8-plex DDI format, all five cAMP derivatives were measured in parallel across five temperatures, yielding distinct apparent thermodynamic fingerprints that are consistent with differences in linker position and chemistry on the adenine ring. A complete five-temperature thermodynamic profile for five compounds with repeated SCK cycles can be acquired within a single day once the conjugates are prepared.

Together, these capabilities make focal molography with DNA-directed immobilization a practical framework for comparative thermodynamic fingerprinting, complementing rather than replacing established calorimetric and refractometric methods.

## Acknowledgments

We thank Daniela Bertinetti and Friedrich W. Herberg (Department of Biochemistry, University of Kassel, Germany) for providing the recombinant PKA-R protein and for its expression and purification, as well as for valuable scientific discussions and critical reading of the manuscript. We gratefully acknowledge the lino Biotech team for their support with laboratory work and data processing, as well as M. Hansch for excellent technical assistance. We thank Marco Campanini (lino Biotech AG) for the software implementation of the refractometric channel.

## Conflict of Interest

Andreas Frutiger and Simona Notova are part of lino Biotech AG, a company involved in the commercialization of focal molography. John Oehninger was affiliated with lino Biotech AG during his internship but is no longer associated with the company.

## Author contributions

Investigation: AF, JO (focal molography experiments); SN (sensor chip preparation and cAMP derivative conjugation). Formal analysis: JO, AF. Visualization: JO, AF. Writing – original draft: JO, AF. Writing – review & editing: JO, AF. (Initials: JO = John Oehninger, SN = Simona Notova, AF = Andreas Frutiger.)

## Artificial intelligence tools statement

Anthropic Claude Opus 4.8, Anthropic Claude Sonnet 4.6, and OpenAI GPT-5.5 were used to assist with data evaluation and manuscript writing. All AI-assisted output was developed, overseen, reviewed, and corrected by the human authors, who take full responsibility for the final content of the manuscript.

## Funding

This work was funded by lino Biotech AG. The funder provided support in the form of salaries for authors AF, SN, and JO (employees or former intern of lino Biotech AG), but had no role in study design, data collection and analysis, decision to publish, or preparation of the manuscript.

## Data availability

All raw data, processed results, and analysis scripts supporting the findings of this study are provided as Supporting Information (supporting information.zip).

## Ethics statement

Human serum preparations used in this study (Serum 1, lot HD2403001, Seqens; Serum 2, lot U1123331304, Sigma-Aldrich) were purchased as commercially sourced, anonymized materials and were not collected specifically for this research. No patient data, personal information, or biological samples obtained from identifiable individuals were used. Accordingly, institutional ethical approval was not required for this study.

## Supporting information

**S1 File. Supporting Information: supplementary figures S1–S8 and supplementary tables S1–S4**. A single combined PDF containing all supporting material referenced in the main text: chemical structures of the five cAMP analogs (S1 Fig); MACS^®^ Matchmaker focal molography setup and flow-chamber dimensions (S2 Fig); temperature-ramp validation on the MACS^®^ Matchmaker (S3 Fig); specificity and control experiments, including positive/negative controls and serum non-specific-binding blanks (S4 Fig); all-16-sensor 1:1 Langmuir fits in buffer and 50% human serum at 20 °C (S5 Fig); Callisto Pre-series prototype instrumentation (S6 Fig); complete DDI-SCK raw sensorgrams and per-sensor fits across all five temperatures (S7 Fig); trypsin-EDTA surface regeneration (S8 Fig); and the supporting tables: Callisto injection sequence (S1 Table), triplicate kinetic summary (S2 Table), mean kinetics with confidence intervals (S3 Table), and DDI immobilization levels (S4 Table).

## Notes

### Competing Interest Statement

I have read the journal's policy and the authors of this manuscript have the following competing interests: Andreas Frutiger and Simona Notova are employees of lino Biotech AG, a company involved in the commercialization of focal molography. John Oehninger was affiliated with lino Biotech AG during an internship but is no longer associated with the company. There are no patents, products in development, or marketed products to declare beyond this. This does not alter our adherence to PLOS ONE policies on sharing data and materials.

